# Assessing the Heterogeneity of Cardiac Non-myocytes and the Effect of Cell Culture with Integrative Single Cell Analysis

**DOI:** 10.1101/2020.03.04.975177

**Authors:** Brian S. Iskra, Logan Davis, Henry E. Miller, Yu-Chiao Chiu, Alexander R. Bishop, Yidong Chen, Gregory J. Aune

## Abstract

Cardiac non-myocytes comprise a diverse and crucial cell population in the heart that plays dynamic roles in cardiac wound healing and growth. Non-myocytes broadly fall into four cell types: endothelium, fibroblasts, leukocytes, and pericytes. Here we characterize the diversity of the non-myocytes *in vivo* and *in vitro* using mass cytometry. By leveraging single-cell RNA sequencing we inform the design of a mass cytometry panel. To aid in annotation of the mass cytometry datasets, we utilize data integration with a neural network. We introduce approximately 460,000∼ single cell proteomes of non-myocytes as well as 5,000∼ CD31 negative single cell transcriptomes. Using our data, as well as previously reported datasets, we characterize cardiac non-myocytes with high depth in six mice, characterizing novel surface markers (CD9, CD200, Notch3, and FolR2). Further, we find that extended cell culture promotes the proliferation of CD45+CD11b+FolR2+IAIE- myeloid cells in addition to fibroblasts.

## Introduction

In recent years, non-myocyte biology has grown to become an important field within molecular cardiology. High quality atlases of non-myocyte abundance (Pinto et al., 2016) and heterogeneity (Farbehi et al., 2019; Skelly et al., 2018) have been published, as well as fascinating studies that have demonstrated the vital importance of non-myocytes on modulating cardiac function, including cardiac growth (Maliken Bryan D. et al., 2018) and maintenance of the cardiac conduction system (Hulsmans et al., 2017). With those recent great advancements in mind, it has become paramount to understand how non-myocytes interact with each other and respond to microenvironmental cues. Cardiac tissue homeostasis is dependent upon non-myocyte signaling and thus, represents a major target for future drug design. Yet many questions still remain unanswered and there are a great number of challenges in studying cardiac non-myocytes.

Cardiac non-myocytes occupy a small percentage of the total volume of the heart, yet compose approximately 70% of all cells in the heart (Pinto et al., 2016). Studying the heart with bulk methodologies obscures biological changes in non-myocytes, impeding detailed and specific analysis of this population. Recent advances within the field have relied upon tissue or cell-specific Cre drivers (He et al., 2017), which enable specific recombination in a small subset of cells. Further, single cell analysis has also been extraordinarily effective in understanding the single cell biology of non-myocytes. However, both of these methodologies are not amenable to high-throughput analysis of many samples and are quite costly, impeding research that utilizes them. Cell culture is better suited to the aforementioned approaches, but there are concerns as to the *in vivo* generalization of *in vitro* models. To address these issues, we present a mass cytometry (CytOF) based characterization of murine cardiac non-myocytes *in vivo* and *in vitro*.

Advances in single cell analysis have enabled broad surveys of high numbers of cells with respect to transcriptomes and proteomes. Various methods for data integration have been developed as well, enabling researchers to perform analyses that incorporate observations from previous studies. Essentially, we are now able to measure different –omic methods under different conditions and integrate them to extract out a coarse-grained understanding of biology at the single cell level. To this end, we employ a sparse autoencoder for clustering, imputing, and embedding (SAUCIE) (Amodio et al., 2019). This deep neural network architecture allows for processing many measurements, as well as the capacity to integrate measurements that have a complex non-linear relationship, such as the expression of transcripts and proteins. Using SAUCIE, we are able to perform data integration to effectively study cellular heterogeneity across multiple modalities and conditions.

In this study, we present the first single cell proteomic measurements of non-myocytes using mass cytometry (CytOF), consisting of a barcoded experiment including a single cell suspensions from six mouse hearts, a pooled reference sample, and four *in vitro* conditions. A total of approximately 460,000 high-confidence single cell proteomes were surveyed in this study. We also present a novel scRNAseq dataset sampled with a rapid preparation method in which CD31+ cells were depleted. All the aforementioned datasets as well, as those described in Skelly and Farbehi were then integrated together using SAUCIE, for a total of approximately 500,000 single cell measurements (Farbehi et al., 2019; Skelly et al., 2018). With this integrated dataset, we identify major cell types –within each dataset and then characterize the heterogeneity of the cells separately, enabling cross comparison. This resource provides novel insights into non-myocyte biology and provides an essential characterization of how cell culture impacts non-myocyte biology.

## Results

### Development of a CytOF panel to assess cardiac non-myocytes

Our experimental design choices were informed by the goal of assessing both *in vivo* and *in vitro* samples. Within cell culture, endothelial cells are typically lost, so we prioritized characterization of the CD31 negative cell population. In order to develop an optimal CytOF panel to assess cardiac non-myocyte heterogeneity, we performed single cell RNA sequencing (scRNAseq) on single cell suspensions derived from the ventricular tissue of a male and female mouse (Fig 1a). Ventricular tissue was digested using a collagenase IV and dispase II solution, after which the resulting suspensions were depleted of endothelial cells using anti-mouse CD31 magnetic beads and a MACS column. We rationalized using MACS to prepare the single cell suspensions would be more time efficient than sorting using FACS, thereby minimizing transcript degradation. The resulting cells were then loaded into a 10X Genomics Chromium capture chip and subsequently sequenced using a NextSeq 500, achieving a read depth of approximately 105,000 reads/cell.

**Figure 1.**
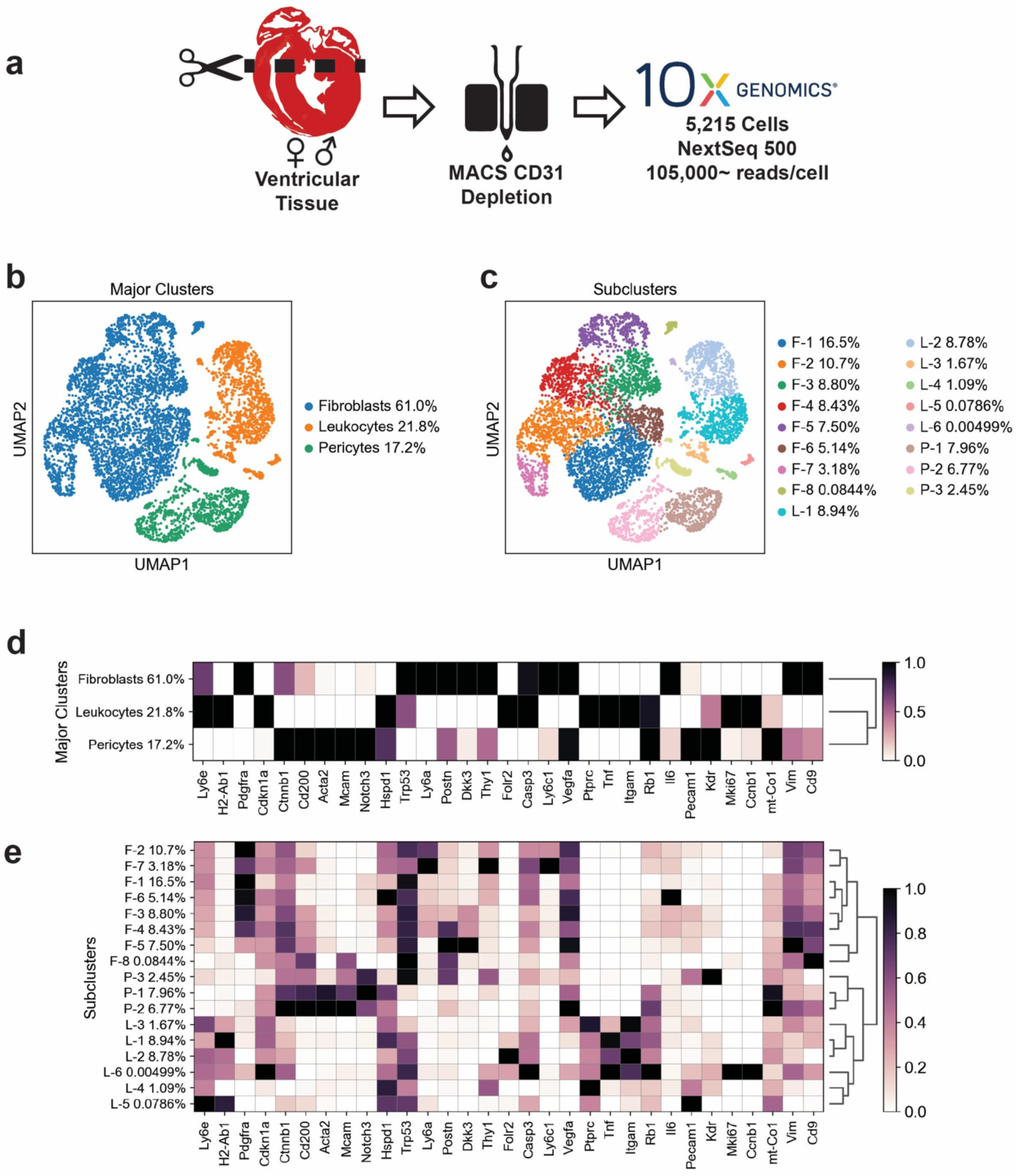
scRNAseq guides CytOF panel design through identification of highly variable genes in cardiac non-myocytes **a)** Experimental workflow and description for scRNAseq experiment. **b)** UMAP embedding of scRNAseq with major cell type labels. **c)** UMAP embedding of scRNAseq data with Leiden community detection-identified subclusters labeled with major cell type and percent composition. **d)** Matrix plot binned by major clusters with panel genes displayed as standard-scaled expression. Dendrogram is defined on the principal component space. **e)** Matrix plot binned by major clusters with panel genes displayed as standard-scaled expression. Dendrogram is defined on the principal component

With this approach, we were able to identify 3 major cell types: fibroblasts, leukocytes, and pericytes (Fig 1b). Of these cell types, we found 8 distinct subpopulations of fibroblasts, 6 distinct subpopulations of leukocytes, and 3 distinct populations of pericytes (Fig 1c). Using this data, we explored the differentially expressed genes found in the major cell types and cross-referenced them with antibody catalogues in order to identify candidate markers. We wanted to characterize cell cycle progression, as it is a major source of cellular heterogeneity. Trp53 (P53), Cdkn1a (P21), mt-Co1 (MTCO1), Hspd1 (HSP60), Tnf (TNFa), and Il6 (IL6) were selected due to their association with cellular stress responses. We reasoned that the expression of Dkk3 would be mirrored by beta-Catenin activity. With scRNAseq, we found that our panel of candidate genes was able to identify a unique gene expression signature associated with the major cell types we identified (Fig 1d) and elucidate aspects of the heterogeneity of subpopulations of non-myocytes (Fig 1e). Some of the markers we identified, namely Cd9, Cd200, Notch3, and Folr2 were never previously characterized in detail on cardiac non-myocytes.

Using CytOF we were able to capture a greater number of cells, conduct more complex experimental designs, and minimize marker detection variability with barcoding. To maximize the efficacy of CytOF, we used a data driven approach to identify candidate genes that would be studied with single cell proteomics, such to recapitulate the power of scRNAseq in cell type classification. While our scRNAseq approach neglected to include the endothelial cells, we reasoned that they could be labeled with a combination of Pecam1 (CD31) and Kdr (VEGFR2), which have been previously established as robust markers for this cell type.

### Description of CytOF experiments and integrated data analysis approach

In order to obtain single cell suspensions for *in vivo* mass cytometry (CytOF), a single cell suspension was generated from 3 male/female gender littermate pairs of mice (Fig 2a). A day (24 hours) before sacrifice, mice were injected with 10 mg/kg 2-iododeoxyuridine (IdU) to label replicating cells for mass cytometry. Hearts were processed as previously described, except in this case, a CD31 depletion was not performed and a debris removal step was performed (Miltenyi debris removal solution). Following debris removal cells and staining with cisplatin, cells were fixed in 4% formaldehyde and stored in Fluidigm Cell Staining Buffer (CSB) with 10% DMSO. An overdigested reference sample was made by digesting a pool of 10 mouse hearts in a large volume of digestion solution for 1 hour, as opposed to 30 minutes. This allowed us to use a reference sample to evaluate the effect of prolonged digestion on the cellular composition of suspensions.

**Figure 2.**
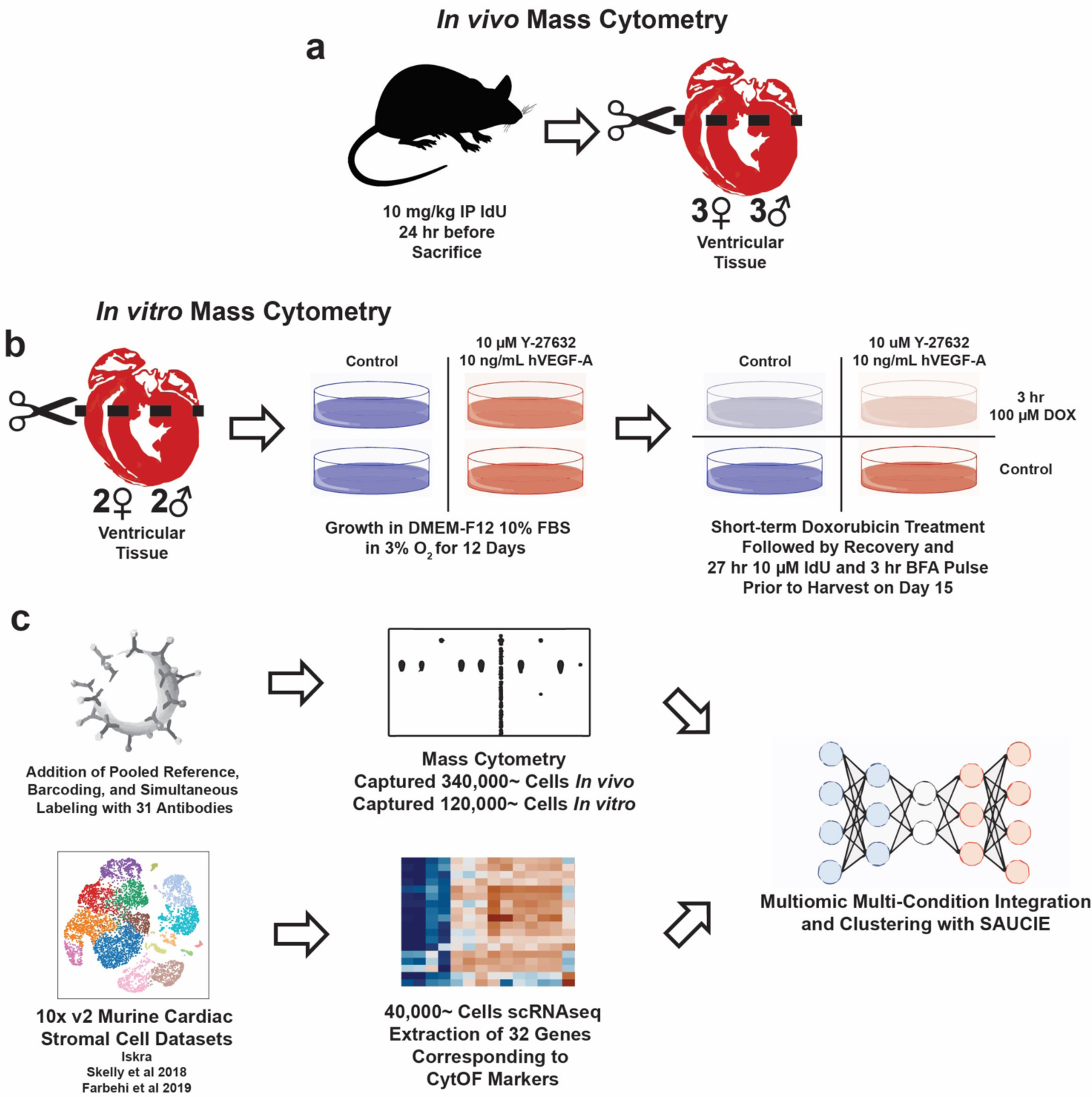
CytOF experimental overview and data merging strategy. **a)** Experimental workflow to generate the *in vivo* CytOF dataset. **b)** Experimental schedule for *in vitro* experiments. Cultures were grown in continuous exposure to DMEM-F12 with (E) and without (V) supplementation of a cocktail of Y-27632 and VEGF followed by 3 hour treatment with doxorubicin (D) or vehicle (V). **c)** Diagram depicting the data integration strategy of our analysis.

Single cell suspensions for *in vitro* samples were derived from 2 female and 2 male mice, which were seeded into 10 cm dishes for growth in physiological normoxia (3% O2) (Fig 2b). Cells were either cultured in DMEM-12 with 10 µM Y-27632 (a Rho-Kinase inhibitor) and 10 ng/mL hVEGF-A (abbreviated as E) or control media (V). Cells were continuously exposed to drug treatments and grown past confluence with media changes every 3 days. At 12 days post seeding, cells were given a 3 hour doxorubicin (DOX) exposure (D) or control media (V), to inhibit cell growth. Media was replaced after treatment and conditions were maintained for 48 hours after exposure, when 10 µM IdU was spiked into culture for 24 hours, followed by a 3 hour brefeldin-A exposure (performed to enhance cytokine staining), and cell harvest. Cells were dissociated by scraping and the addition of TrypLE. Cells were digested for 45 minutes at 37 in TrypLE. Cells were stained with cisplatin, fixed, and stored for staining. All four conditions are referred to as VV, VD, EV, and ED. As evidenced by other investigators, we rationalized hypoxia would enhance cell growth and growing cells in a post confluent state would promote greater culture heterogeneity (Ackers-Johnson Matthew et al., 2016). Y-27632 and VEGF-A were used as a supplement, as they have been previously found to increase the proliferation of CD31+ cells in culture (Joo et al., 2012); additionally, Y-27632 has been demonstrated to inhibit fibroblast-to-myofibroblast transition. Cells were treated with DOX in order to arrest the cell cycle to better identify replicating cells.

Single cell suspensions for mass cytometry were thawed and barcoded to enable simultaneous staining of samples, which allowed us to minimize staining variability between samples and filter out doublets. A suspension of singly-stained polystyrene antibody binding beads was run separately from the sample on the same day to allow us to correct for spillover from antibodies. A listing of antigens stained, clones of the antibodies used, and putative transcriptomic correlate or mapping is provided in Table 1. Raw fcs files were obtained from the Helios and then processed with CATALYST (Chevrier et al., 2018) in order to normalize, compensate, and debarcode data files. Following debarcoding, fcs files were gated in accordance with Fluidigm recommendations using FlowJo in order to remove doublets based on the Helios Guassian gates. After this, an arcsinh5-transformed count matrix was obtained and unrandomized using a ceiling function.

**Table 1.** List of gene symbols, protein marker, antibody clone, metal conjugate, and marker type/description.

| Gene Symbol | Protein | Antibody Clone | Metal | Marker Type |
| --- | --- | --- | --- | --- |
| Ptprc | CD45 | 30-F11 | 89Y | Leukocyte Identity |
| Itgam | CD11b | M1/70 | 143Nd | Myeloid Identity |
| Folr2 | PE-FolR2 | FR-β | Anti-PE<br>145Nd | L-2 Identity |
| H2-Ab1 | IAIE | M5/114.15.2 | 209Bi | L-1 Identity |
| Tnf | Tnfa | MP6-XT22 | 141Pr | Leukocyte Cytokine |
| Pdgfra | CD140a | APA5 | 148Nd | Fibroblast Identity |
| Vim | Vimentin | D21H3 | 154Sm | Fibroblast Identity |
| Thy1 | CD90 (CD90.2) | 30-H12 | 156Gd | F-1/2/6 Identity; P-3 Identity |
| Ly6c1 | Ly6C | HK1.4 | 150Nd | F-2 Identity; Endothelial Identity |
| Ly6a and Ly6e | Ly6AE (Sca-1) | D7 | 169Tm | Fibroblast Identity; Endothelial Identity |
| Postn | Postn | EPR20806 | 170Er | F-3/4/5 Identity; P-3 Identity; Myofibroblast Identity |
| Il6 | Il6 | MP5-20F3 | 167Er | Fibroblast Cytokine |
| Cd9 | CD9 | KMC8 | 158Gd | Fibroblast Identity; Subtype for all Cells |
| Vegfa | VEGF | VG1 | 174Yb | Fibroblast Cytokine |
| Mcam | FitC-CD146 | ME-9F1 | Anti-FitC<br>160Gd | Pericyte Identity; Endothelial Identity |
| Cd200 | APC-CD200 | OX-90 | Anti-APC<br>176Yb | Pericyte Identity; Endothelial Identity |
| Notch3 | Notch3 | AF1308 | 173Yb | Pericyte Identity |
| Acta2 | αSMA | MAB1420 | 164Dy | P-1/2 Identity; Myofibroblast Identity |
| Pecam1 | CD31 | MEC13.3 | 168Er | Endothelial Identity |
| Kdr | VEGFR2 | MAB4432 | 159Tb | Endothelial Identity |
| Casp3 | ActCaspase3<br>(Cleaved Caspase 3) | D3E9 | 142Nd | Apoptosis; Cell Stress Marker |
| Trp53 | P53 | Pab240 | 171Yb | Apoptosis; Cell Stress Marker |
| mt-Co1 | MTCO1 | EPR19628 | 151Eu | Mitochondrial content; mtDNA Encoded |
| Hspd1 | HSP60 | EPR18245-93 | 162Dy | Mitochondrial content; nDNA Encoded |
| Mki67 | Ki67 | B56 | 161Dy | Cell Cycle; S-Phase |
| Rb | pRb (phospho-Retinoblastoma) | D20B12 | 166Er | Cell Cycle; G1-Phase |
| Ccnb1 | CyclinB1 | EPR17060 | 153Eu | Cell Cycle; G2/M-Phase |
| Birc5 | pHistoneH3 (phospho-HistoneH3) | HTA28 | 175Lu | Cell Cycle; M-Phase |
| Cdkn1a | P21 | EPR362 | 149Sm | Cell Cycle; G0-Phase |
| Ctnnb1 | bCatTotal | D10A8 | 147Sm | β-Catenin Cytoskeletal Content |
| Dkk3 | bCatActive | D13A1 | 165Ho | Wnt/β-Catenin Activity |

Data integration with SAUCIE allowed us to take the different CytOF datasets as well as previously published v2 10X Genomics datasets, merge them into a shared embedding space, and generate a reconstructed gene expression value (designated by the suffix ‘_r’). Previously published data from Skelly (containing flow sorted, CD31 depleted homeostasis data) and Farbehi (containing myocardial infarction/MI data) was recounted using CellRanger 3.0.2 and integrated into our dataset without the need for batch correction. Including our dataset, a total quantity of 42,843 single cells transcriptomes were used for this study. Single cell transcriptomes were processed with MAGIC (Dijk et al., 2018) prior to integration with SAUCIE to smooth single cell transcriptome data, which enhanced data integration. Protein count matrices from each experiment were loaded into Scanpy and processed to remove low quality cells in CytOF data, giving us 343,245 cells from the *in vivo* CytOF and 125,109 cells from the *in vitro* CytOF datasets (Fig 2 Supplement 1). Each transformed protein marker was mapped to a transcript (Table 1) and then loaded into a data matrix containing 32 channels (Fig 2c). In the cases of IdU, beta-Catenin activity, and pHistoneH3 expression, we assumed their expression was related to Ki67, Dkk3, and Birc5, based on gene-ontology association. The resulting data matrix was then used to train SAUCIE to map mass cytometry measurements to transcriptomic measurements.

Training data from scRNAseq and *in vitro* experiments was randomly sampled with replacement to prevent overfitting on *in vivo* data. Optimal hyperparameters were found based on recommendations published within the paper describing SAUCIE, in so much that lambda B was chosen to minimize distances between the shared sample latent space without perfectly blending the data and lambda D was set to double the value of lambda C. Analysis performed at different hyperparameters were able to recapitulate the clustering with minor amounts of variability. In addition, we ensured that the latent space was able to recapitulate known minor cell populations, such as lymphocytes (CD45+CD90+; Ptprc+Thy1+), and our clustering analysis was able to recapitulate previous analyses, to ensure biologically meaningful results. We found that integration of MAGIC-preprocessed scRNAseq data facilitated better accuracy of SAUCIE reconstructions and more meaningful clustering results.

### Integrated analysis reveals the proportions and relationships of major cell types across experiments

With SAUCIE, we were able to identify the expected four major cardiac non-myocytes: endothelium, fibroblasts, and pericytes. Cell identity within the SAUCIE embedding (Fig 3a) is related to its position. By examining the expression of markers within the SAUCIE embedding (Fig 3b), we were able to determine the cell composition of each cluster. Throughout all the experiments, we found robust representation of fibroblasts, followed by endothelial cells, leukocytes, and pericytes (Fig 3c). This is interesting, as endothelial cells are the most abundant non-myocyte. Based on this analysis, our knowledge of single cell biology of fibroblasts is greatest, whereas that of pericyte single cell biology is the least.

**Figure 3.**
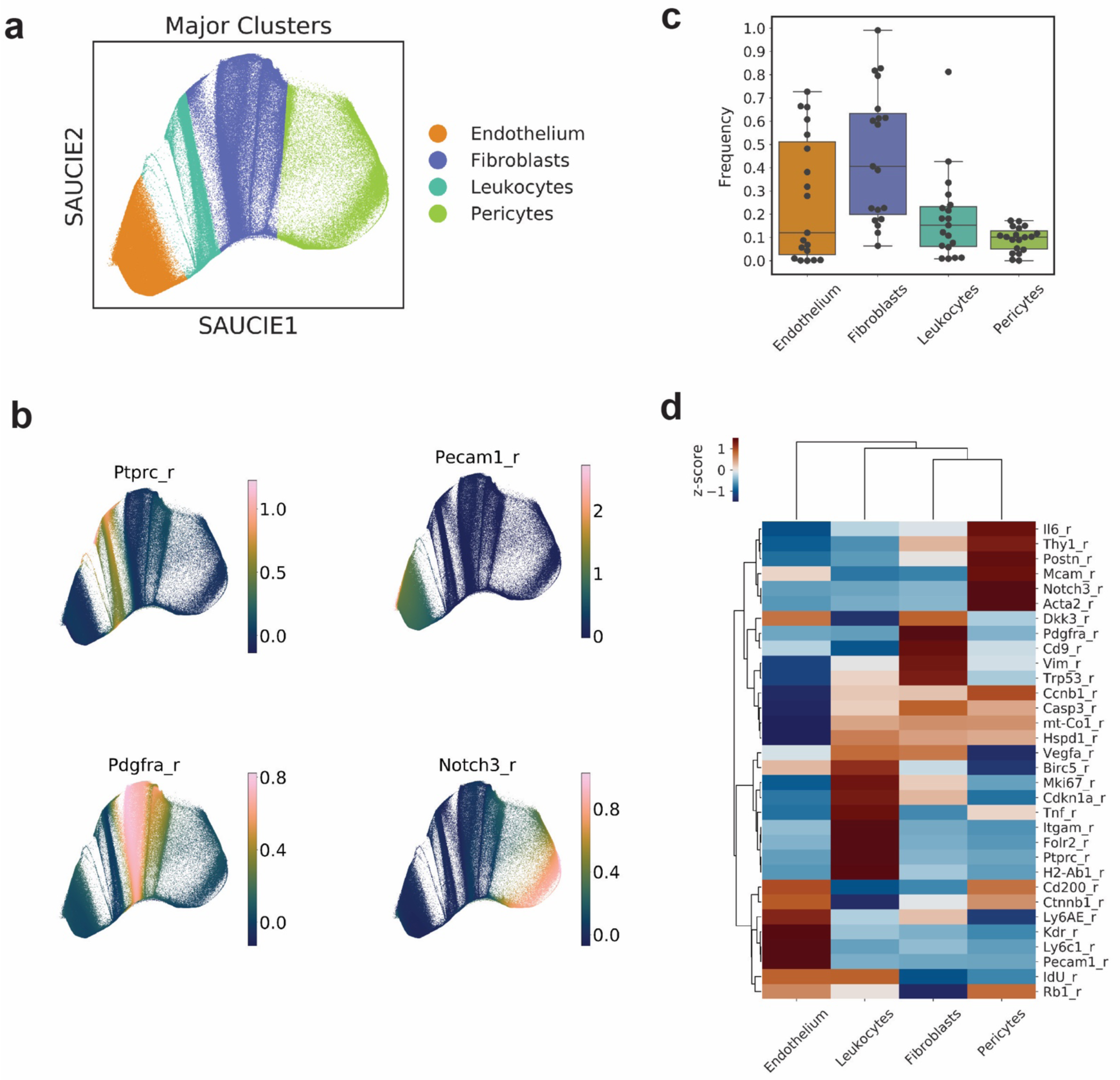
SAUCIE-integration of datasets facilitates identification of four major cell types. **a)** Major clusters identified within SAUCIE clustering analysis plotted on the SAUCIE embedding space. **b)** Feature plots of major markers used to facilitate major cluster identification. **c)** Frequency of major cell types identified across all datasets. **d)** Heatmap of SAUCIE reconstructed gene expression profiles for the major cell types. For boxplots, the box highlights quartiles, whereas the whiskers highlight the data range, and outliers fall off the range.

Further, we found strong concordance between the integrated gene-expression data (Fig 3d) and our scRNAseq data (Fig 1d), with some exception. Chiefly, the expression of inflammatory markers (TNFa and IL6) and mitochondrial markers (MTCO1 and HSP60) were found to be discordant. Surprisingly, P53 and P21 were found to be associated with fibroblasts and leukocytes, respectively. Cell cycle marker expression levels for the major cell types closely followed their scRNAseq expression levels, suggesting that distinct cell types have a tendency to express different cell cycle markers. Also, we found endothelial cells to express low amounts of mitochondrial associated proteins, which is surprising, given the central roles mitochondria play in the endothelial cell biology (Kluge et al., 2013). Consistent with past findings (Kretzschmar et al., 2018), IdU labeling was found to be most robust in endothelial cells and leukocytes.

Next, we examined each modality (*in vivo* CytOF, *in vitro* CytOF, and scRNAseq) by itself, using SAUCIE clusters. *In vivo* samples were found to occupy a distinct section of the SAUCIE embedding space (Fig 4a). The ratios of non-myocytes identified was concordant with Pinto 2016; however, we were able to detect a greater degree of endothelial cell variability (Fig 4b). We believe this is a result of endothelial cells being sensitive to dissociation, a hypothesis that is validated by our overdigested reference sample, where endothelial cells were found to compose about 30% of all identified cells. *In vitro* cells were found to have little overlap with *in vivo* cells (Fig 4c), suggesting a divergence in phenotype induced by cell culture. We also found variability in the proportions of cells identified within *in vitro* CytOF data, likely secondary to culture conditions. The majority of cells found within *in vitro* conditions were fibroblasts (Fig 4d) with variable amounts of leukocytes, dependent on culture conditions.

**Figure 4.**
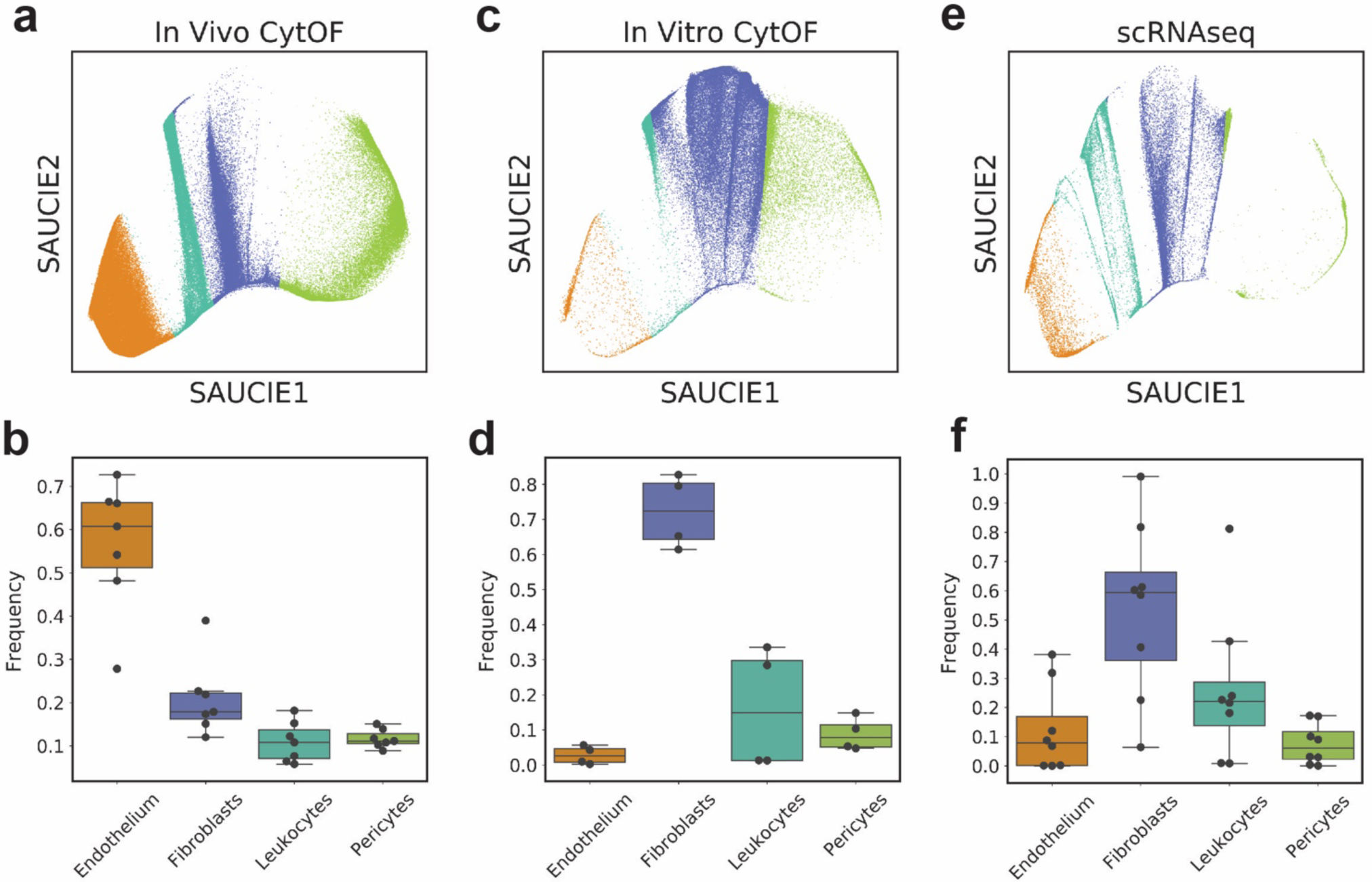
Localization in the SAUCIE embedding space and frequency of cell types by data modality. **a)** Distribution of all *in vivo* data in the SAUCIE embedding space. **b)** Boxplot of frequency of cell types within the *in vivo* data. **c)** Distribution of all *in vitro* data in the SAUCIE embedding space. **d)** Boxplot of frequency of cell types within the *in vitro* data. **e)** Distribution of all scRNAseq data in the SAUCIE embedding space. **f)** Boxplot of frequency of cell types within the scRNAseq data. For boxplots, the box highlights quartiles, whereas the whiskers highlight the data range, and outliers fall off the range.

By examining the SAUCIE embedding space of scRNAseq datasets, we found that the distribution of cells mostly matched *in vivo* cells with some exception (Fig 4e). The scRNAseq datasets contained data from Farbehi 2019, which profiled cells derived from infarcted hearts. Therefore, some of the cells found to overlap with *in vitro* data, came from those datasets, indicating that *in vitro* conditions may represent injury conditions. In particular, we expect that some leukocytes, fibroblasts, as well as a cell type ambiguously marked by both fibroblasts and pericyte (likely myofibroblasts) belonging *in vitro* are representative of injury-associated cells. For this reason, we believe that *in vitro* non-myocytes are a powerful model to study cardiac injury response.

With this analysis, we were able to profile many datasets at once and identify the proportions of endothelium, fibroblasts, leukocytes, and pericytes with a shared label. We found that *in vitro* cells likely represent cell types present during cardiac injury recovery.

### Comparative analysis of cellular heterogeneity in CytOF datasets

In order to identify the cell types within each dataset with greater accuracy, we performed clustering analysis with Louvain clustering as implemented in scanpy. It is important to state here that *in vivo* and *in vitro* cells are of different sizes, and that is reflected within our datasets (Supplementary Figure 1). We found that cells *in vitro* had greater median protein content and a greater median diversity of genes expressed than cells found *in vivo*. For these reasons, we did not integrate the *in vivo* and *in vitro* datasets outside of the SAUCIE analysis. Due to the low quantity of endothelium found within our *in vitro* experiments, we excluded them from the analysis of the *in vitro* and *in vivo* datasets, as we were not able to confidently sample them for comparisons. For this analysis, 45,000 cells were subset from each CytOF dataset and analyzed separately.

First, we examined the expression of CytOF protein markers on a per cell basis using the SAUCIE Major Clusters labels. Cells found *in vivo* were well defined based on their expression patterns for major markers (Fig 5a). Fibroblasts were found to express the highest amounts of CD140a, Vimentin, and CD9. Interestingly, we found that fibroblasts expressed CD45 and IAIE, suggestive of a potential lineage relationship with cardiac leukocytes. Indeed, other groups have observed the expression of CD45 and IAIE on fibroblasts, and they have been referred to them as myeloid fibroblasts (Cieslik et al., 2015; Trial et al., 2017) or fibrocytes (Skelly et al., 2018). Using CytOF, which can detect the presence or absence of markers with greater accuracy than fluorescence based flow cytometry, we observed an unprecedented amount of these surface markers on fibroblasts. Other surface markers of fibroblasts, such as CD90, Ly6AE, and Ly6C were detected on fibroblasts with high amounts of variability. Leukocytes were found to express CD45, CD11b, IAIE, and FolR2 highly, with FolR2 expression being the most variable surface marker. In addition, we found that leukocytes expressed the highest amount of pHistoneH3, suggesting a greater amount of mitotic cells within this population of cells. Pericytes were marked high CD90, CD146, CD200, Notch3, bCatTotal, and aSMA expression, with Notch3 expression being the most specific marker of pericytes. Interestingly, bCatActive was found to be expressed primarily on pericytes.

**Figure 5.**
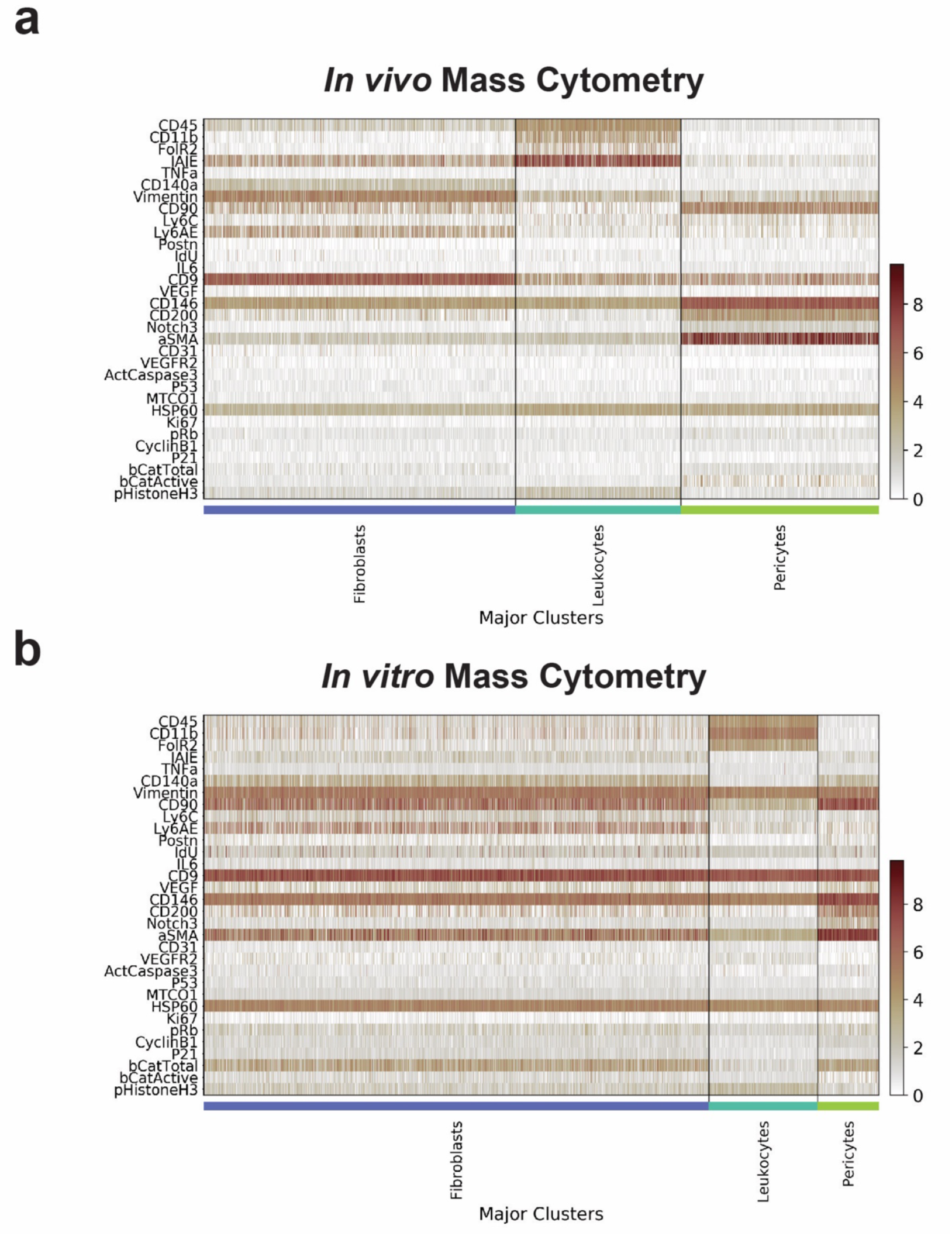
Comparative examination of single cell expression profiles from CytOF datasets. **a)** Single cell heatmap of cells from the *in vivo* CytOF dataset. Expression is represented as hyperbolic arcsin factor 5 transformed counts. **b)** Single cell heatmap of cells from the *in vitro* CytOF dataset. Expression is represented as hyperbolic arcsin factor 5 transformed counts.

*In vitro* cells were found to be overall less variable than *in vivo* cells with respect to marker expression (Fig 5b). Vimentin and CD9 expression was found to be ubiquitous, demonstrating that cells grown *in vitro* have a fibroblastic phenotype. The defining cell markers for each cell type were expressed in a more ambiguous fashion. Similar to our *in vivo* observations, fibroblasts expressed vimentin, CD9, CD140a, Ly6AE, and Ly6C. However, fibroblasts had increased aSMA and CD90 expression, which is characteristic of fibroblast activation. Further, there was an increase in IdU expression in the absence of increased expression of other cell cycle markers, confirming our previous observations that cell cycle markers have a cell type-specific bias. Leukocytes were found to be uniformly expressing CD45, CD11b, and FolR2, whereas IAIE was found to be absent, which was very different than what is found under *in vivo* conditions. Again, we found that pHistoneH3 expression primarily labeled leukocytes. Pericytes were harder to distinguish from fibroblasts, with CD146, CD200, and Notch3 being positive markers and Ly6AE being a negative marker. aSMA was found to be less variable in this population. It is likely that mature myofibroblasts were identified as pericytes within this analysis, given the similarity of the two cell types.

Louvain clustering was able to identify 18 unique clusters within the *in vivo* dataset (Fig 6a) and 15 unique clusters (Fig 6b) in the *in vitro* dataset, indicative that *in vitro* culture diminishes cellular heterogeneity of cardiac non-myocytes. SAUCIE-identified major cell types labeled distinct clusters in the *in vivo* data (Fig 6c), whereas within the *in vitro* dataset (Fig 6d), cells were more ambiguously labeled. The mixing of labels within *in vitro* clusters suggests a phenotypic shift of leukocytes to a more fibroblastic cell type (myeloid-to-fibroblast transition) as well as fibroblast-to-myofibroblast transition occurring within culture. Evaluating the intersample heterogeneity of the *in vivo* samples demonstrated high amounts of biological variability (Fig 6e). Further, as discussed in the previous section, we found stark differences in *in vitro* sample heterogeneity associated with cell culture conditions (Fig 6f). Treatment with the E cocktail diminished the presence of clusters 0, 1, 5, and 10 and enriched for the presence of clusters 2, 3, 4, 6, 7, 8, 9, 11. DOX treatment was associated with diminishment of clusters 10 and 13, while cluster 12 was found to be enriched.

**Figure 6.**
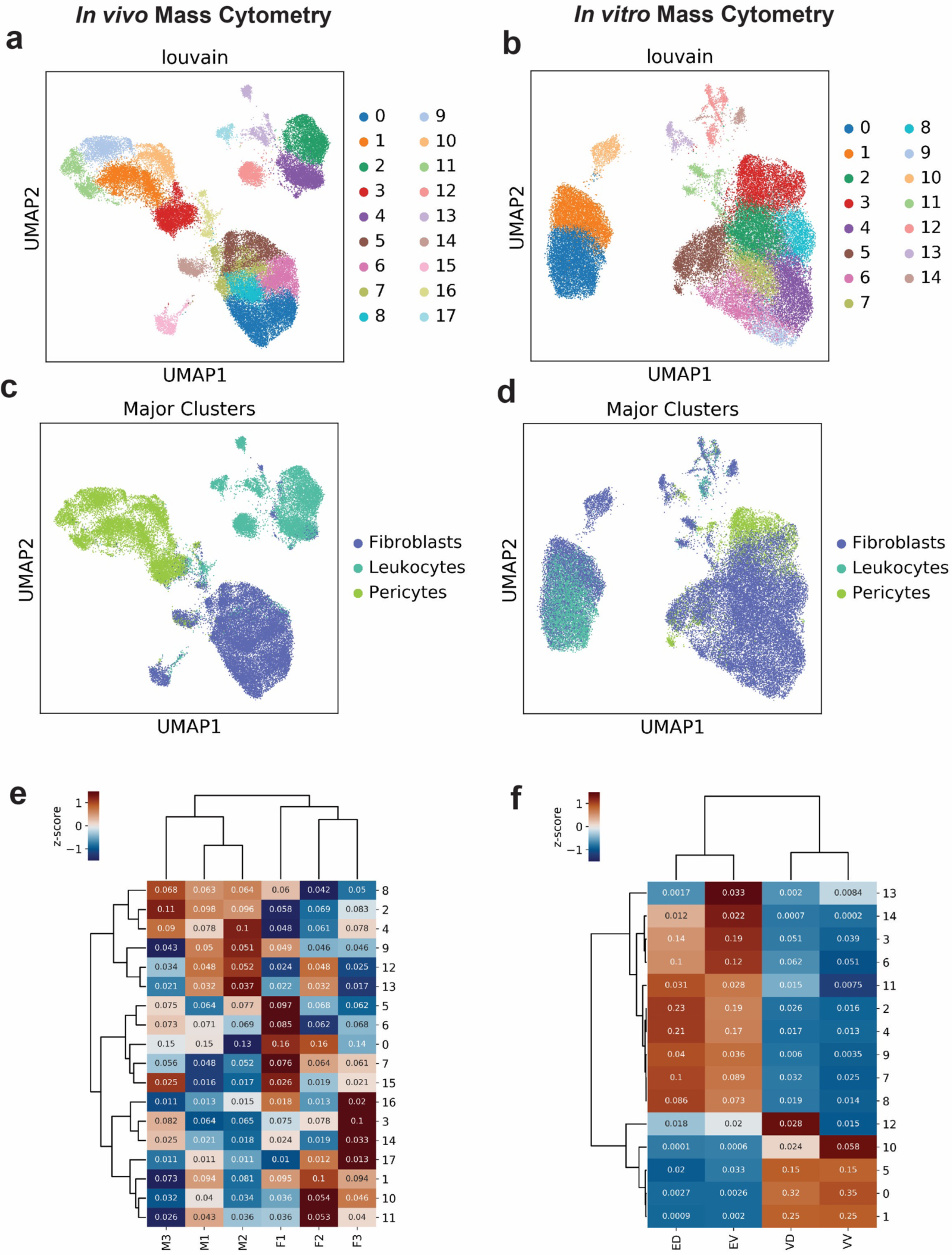
Comparative analysis of unsupervised clustering results and intersample heterogeneity. **a)** Louvain community detection clusters plotted on UMAP embedding from *in vivo* data subset. **b)** Louvain community detection clusters plotted on UMAP embedding from *in vitro* data subset. **c)** SAUCIE Major Clusters plotted on UMAP embedding from *in vivo* data subset. **d)** SAUCIE Major Clusters plotted on UMAP embedding from *in vitro* data subset. **e)** Hierarchically clustered heatmap of *in vivo* cluster abundances by sample. Dendrogram was determined based on the data frame of cluster abundance values found in each sample. Annotation contains the frequency of each cluster. **f)** Hierarchically clustered heatmap of *in vitro* cluster abundances by sample. Dendrogram was determined based on the data frame of cluster abundance values found in each sample. Annotation contains the frequency of each cluster.

Next, we examined the expression profiles of the clusters identified in each modality. Within the *in vivo* experiment, we found high amounts of heterogeneity (Fig 7a). Notably, we found lymphocytes (cluster 17), putative Dkk3+ fibroblasts (cluster 14), putative endothelial progenitor cells or endothelial-to-mesenchymal-transition (EMT) associated cells (cluster 15), as well as an apoptotic cell type without distinguishable surface markers but high expression of secreted molecules (cluster 16). Fibroblasts were represented by clusters 0, 4, 5, 7, 6, and 8, with heterogeneity resulting from Ly6AE^hi^ cells (cluster 0, 6, and 8), CD90^hi^ cells (clusters 0 and 7), and CD45^hi^IAIE^hi^ cells (cluster 6). Aside from the lymphocytes, leukocytes were identified in clusters 2, 4, 12, and 13, with major sources of heterogeneity resulting from CD11b^-^P21^-^ pHistoneH3^+^ cells (cluster 12), FolR2^+^ cells (cluster 2), and Ly6C^hi^IAIE^-^IdU^+^ cells (cluster 13). Pericytes were represented by clusters 1, 3, 9, 10, and 11, with heterogeneity resulting from Ly6AE^hi^Ly6C^hi^ cells (cluster 10), aSMA^-^ cells (cluster 3), and pRb^hi^MTCO1^hi^HSP60^hi^ cells (clusters 9 and 11).

**Figure 7.**
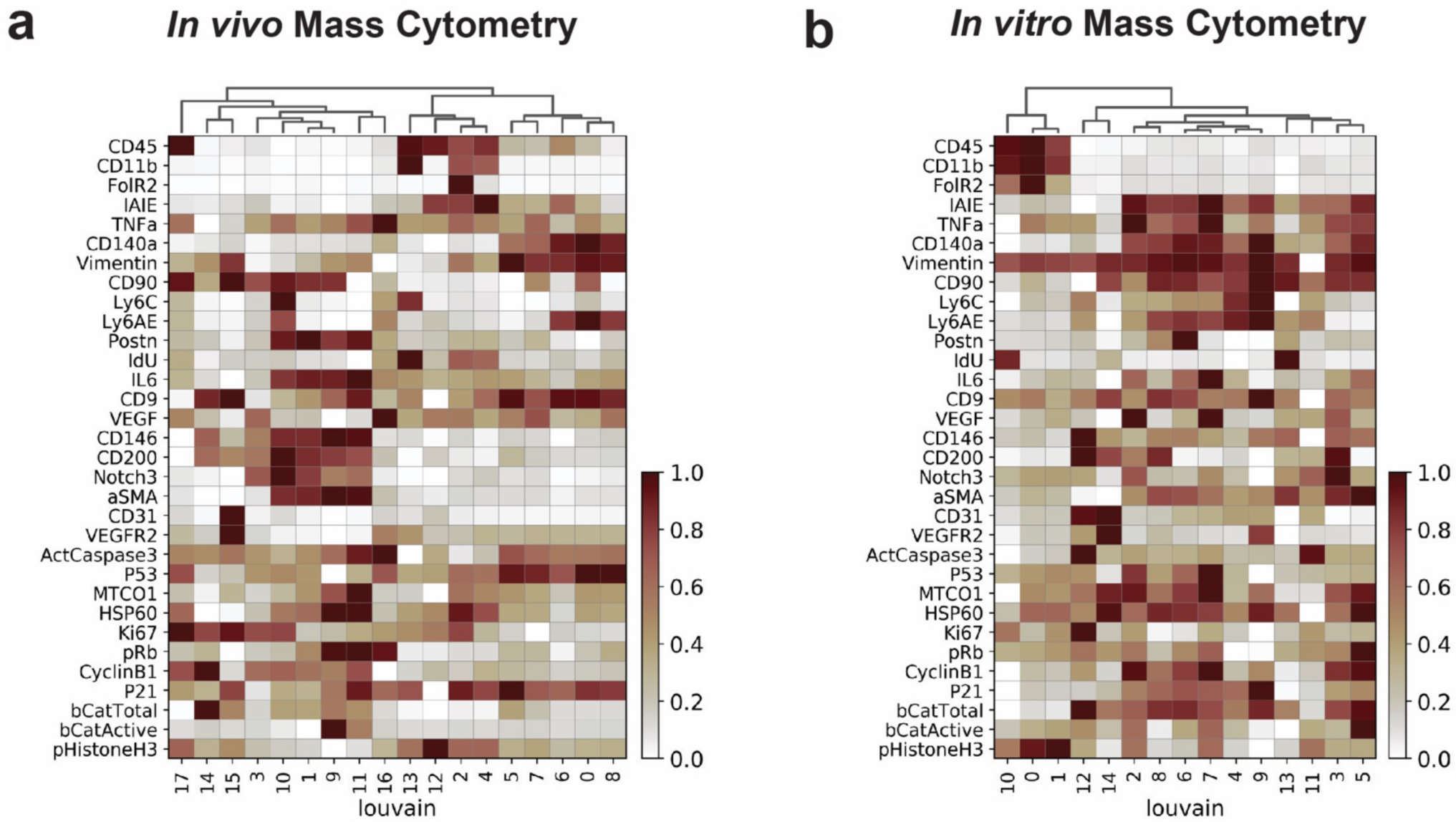
Comparative analysis of cluster expression profiles. **a)** Heatmap single cell expression profiles binned by Louvain community detection clusters with standard scale expression of markers from *in vivo* data subset. Dendrogram based on protein expression values. **b)** Heatmap single cell expression profiles binned by Louvain community detection clusters with standard scale expression of markers from *in vitro* data subset. Dendrogram based on protein expression values.

Within the *in vitro* data (Fig 7b), cells were more monolithic and vimentin expression was found to be uniform, with the exception of cluster 11, which is likely an apopotic cell type similar to cluster 16 within the *in vivo* data, with ambiguous identity but lacking the expression of secreted molecules. Interestingly, clusters 12 and 14 identified a putative endothelial progenitor cell or an EMT associated cell with variable expression of VEGFR2, Ki67, MTCO1, HSP60, and bCatTotal. Fibroblasts were found in clusters 2, 4, 6, 7, 8, 9, and 13 with Postn+ cells identified in cluster 6, IL6^+^VEGF^+^P53^+^MTCO1^hi^CyclinB1^+^ cells found in clusters 2 and 7, Ly6AE^+^Ly6C^+^ cells found in clusters 4 and 9, and IdU^+^ cells found in cluster 13.. The co-expression of markers in clusters 2 and 7 suggest a P53-dependent cytokine secretion regulatory network exists in fibroblasts. In addition, all fibroblasts were found to express high quantities of TNFa. Leukocytes were found in clusters 0, 1, and 10, with 10 having the greatest IdU expression, intermediate FolR2 expression, and least pHistoneH3 expression, whereas 1 and 10 had the highest expression of pHistoneH3 and FolR2 being most highly expressed on cluster 0. Pericytes were found in clusters 3 and 5, which differed in bCatActive, MTCO1, HSP60, CD200, and Notch3 expression.

We were able to identify differentially abundant clusters within the *in vitro* experiment. With the expression profiles in mind, we found that treatment with the E cocktail promoted the abundance of fibroblasts (2, 3, 4, 6, 7, 8, 9, 11, 14) as well as putative endothelial progenitor cells (cluster 14) while the abundance of leukocyte (0, 1, 10) and pericyte (5) clusters was found to be diminished. Treatment with DOX diminished the presence of clusters 10 and 13, both of which correspond to an IdU^+^ leukocytes and fibroblasts respectively. Interestingly, expansion of cluster 12, an endothelial progenitor-like cell was associated with DOX treatment. The observations regarding the heterogeneity of cells detected with the E cocktail are supported by experiments reported by other groups using bulk technologies (Omelchenko et al., 2002), as Y-27632 was found to inhibit fibroblast activation. In this context, we hypothesize that cluster 5 is a mature myofibroblast, which has a similar enough protein expression profile as pericytes to be labeled as such. Cluster 3, which we also identified to be a pericyte most likely reflects some intermediate state between fibroblast differentiation and a mature myofibroblast. Surprisingly, neither cluster 3 nor 5 had high expression of Postn, despite expressing the greatest amount of aSMA. This observation suggests that Postn upregulation is an intermediate step between fibroblast differentiation and mature myofibroblast commitment. Further, it is likely that the presence of leukocytes is dependent upon fibroblast and myofibroblast commitment, as both cell types are highly secretory. However, we did not observe a possible causative cytokine that would promote leukocyte signaling.

We were unable to observe many consistent differences between mouse sex within the Louvain clusters (Fig 6e). It is apparent that murine cardiac non-myocytes have a high degree of biological variability between individual mice. It is likely that the inclusive of many signaling related proteins in our clustering may have obscured the detection of differentially abundant populations. With that said, the greatest signal for differentially abundant populations of cells came from clusters 2 and 4, which indicated a potential increase in the abundance of myeloid cells associated with male sex. Further, upon examination of the Major Clusters detected by SAUCIE, we found a statistically significant two-fold increase in the number of leukocytes in male mouse hearts as well as a statistically significant decrease in the number of endothelial cells in male mouse hearts (Figure 6 Supplement 1).

In summary, our comparative analysis found diminished subpopulation heterogeneity of the previously identified major cell types within *in vitro* conditions. Cell cycle markers were found to be biased in expression based on specific cell types. For instance, leukocytes had the greatest pHistoneH3 expression, pericytes had the greatest Cyclin B1 expression, and pRb, IdU, as well as Ki67 abundance did not vary together. In addition, there was evidence for P53-modulated secretory networks present within fibroblasts. We also found that cells grown *in vitro* adopt a fibroblastic phenotype, with secondary phenotypes corresponding to leukocyte or pericytes being present. Also, a putative endothelial progenitor cell was identified within the *in vitro* and *in vivo* experiments based on expression of fibroblast markers in addition to CD31 and VEGFR2 expression. We also identified that endothelial cell content as well as leukocyte content are significantly different in male versus female mice. Further, we found that *in vitro* culture, in the absence of the E cocktail, CD45^+^CD11b^+^FolR2^+^ leukocytes proliferated, suggestive that cell culture systems can be useful to understand cardiac myeloid cell biology.

## Discussion

Here we present the first ever mass cytometry datasets of cardiac non-myocytes *in vivo* and *in vitro*. Using a data driven approach, we identified markers for our CytOF panel as well as characterized novel markers for cardiac fibroblasts and pericytes. This strategy is scalable to other systems and provides a basis for rational CytOF panel design. The power of high-throughput, high-dimensional measurements is to reduce bias in data acquisition and analysis. Designing CytOF panels solely based on field consensus diminishes some of this power. scRNAseq is an effective technology to develop panels around, as it provides a survey of transcripts, which are not as deeply affected by technical biases as antibodies. In order to increase experimental repeatability, the selection of commonly used clones for specific antigens is important. While this is not always possible for less commonly studied markers, having a list of candidate genes generated from high quality scRNAseq data, greatly facilitates antibody selection. Ultimately, experimental findings require validation and having multiple measurements with scRNAseq and comparing them to measurements made with CytOF is an important part of rigorous experimental design.

Using mass cytometry, we were able to sample many conditions at once while minimizing technical variation and the detection of doublets (Zunder et al., 2015), enabling rigorous comparisons between samples. With this approach, we were able to provide novel insight into cardiac non-myocyte heterogeneity, as well as sex differences *in vivo*. With the higher dimensionality of our measurements, we were able to identify a putative endothelial progenitor or an EMT-associated cell that exists within both *in vivo* and *in vitro* datasets. Further, we identified four novel surface markers, CD9, CD200, Notch3, and FolR2, in the context of cardiac non-myocyte heterogeneity. We found that CD9 is a robust marker of fibroblasts that is similar in expression as vimentin, whereas CD200 and Notch3 mark pericytes, and FolR2 marks a major subset of myeloid cells. This highlights interesting biology for investigation, as these markers likely have functional roles within each of cell type.

Integrated analysis is able to define cell populations across technologies and in distinct experimental systems (*in vivo* vs *in vitro*), as well as incorporate observations from other research groups. For this reason, data integration with SAUCIE can ensure a level of concordance between observations that can be expressed quantitatively and qualitatively. This work provides a basis for a stable concept of four major cell types that exist within 18 different datasets. In addition, more specific differences within the datasets can be interrogated with SAUCIE to identify transcriptomic associations with variations in protein measurements. This is a sustainable strategy to repurpose costly single cell data acquisition methods and leverage the knowledge obtained in previous studies effectively.

With data integration, we are able to effectively understand how novel data can be used to update our understanding of non-myocytes. For instance, with data integration we can easily appreciate that single cell studies have focused on fibroblasts and leukocytes, while endothelial cells have been neglected. Future single cell inquiry can focus on expanding our knowledge of endothelial cells with more effective single cell dissociation methods, which may preserve endothelial cell integrity better. In our experience, endothelial cells are sensitive to dissociation and are difficult to study even with single cell analysis. Through the formulation of better methods to prepare single cell suspension, we can more effectively understand how the most abundant non-myocyte affects cardiac function and contributes to disease pathologies.

In addition, we were able to discern the impact of cell culture on cardiac non-myocytes and identify some bounds upon which cell culture experiments can generalize to *in vivo* studies. Overwhelmingly, we found that *in vivo* and *in vitro* conditions are very distinct, but it is likely that *in vitro* conditions reflect cells found within typical wound healing processes in the heart. This is an unsurprising finding; however, the identification of CD45^+^CD11b^+^FolR2^+^ proliferating in culture is potentially an exciting observation to further understand cardiac myeloid cell biology. Myeloid cells are studied *in vivo* as they are very sensitive to microenvironmental cues and lose their *in vivo* characteristics in culture rapidly (Janssen et al., 2016). The bed of fibroblasts present in overconfluent culture can act as a substitute for the microenvironment, possibly rendering more *in vivo*-like myeloid cells that can be used to investigate more mechanistic questions. We’ve found (data not shown) that in 18 days of culture, the number of leukocytes can expand to over 50% of the culture content, greatly increasing the number of cells that can be assayed.

In summary, we have provided novel datasets with over 400,000 single cell proteomes and 5,000 single cell transcriptomes, a powerful analysis approach that integrates previous observations from other investigators and technologies, as well as a high-dimensional proteomic characterization of non-myocytes with novel markers *in vivo* and *in vitro*. We hope that this resource and the discussion of its findings can benefit the molecular cardiology community and spur further investigation into cardiac wound healing mechanisms.

## Materials and Methods

### Key Resource Table

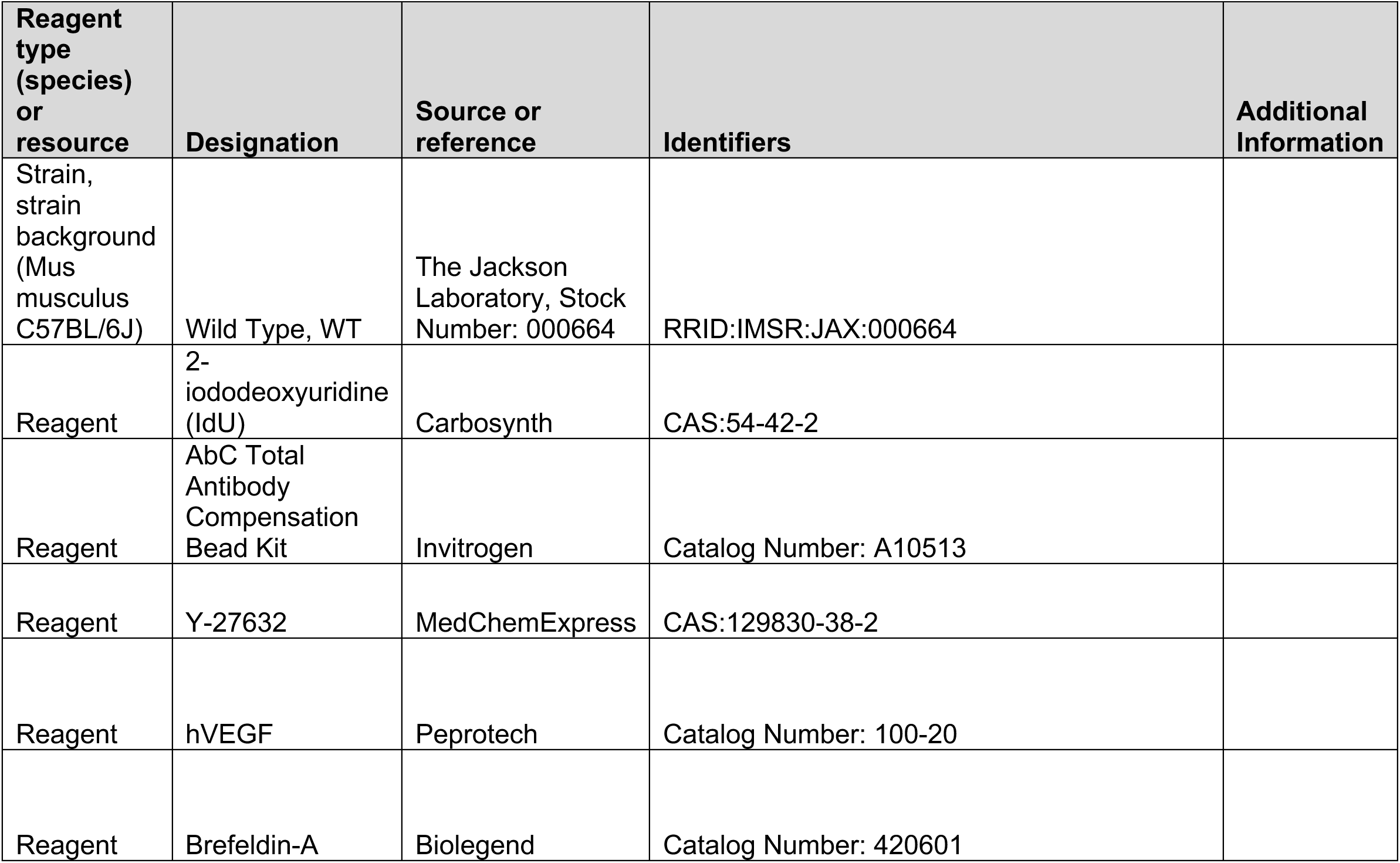
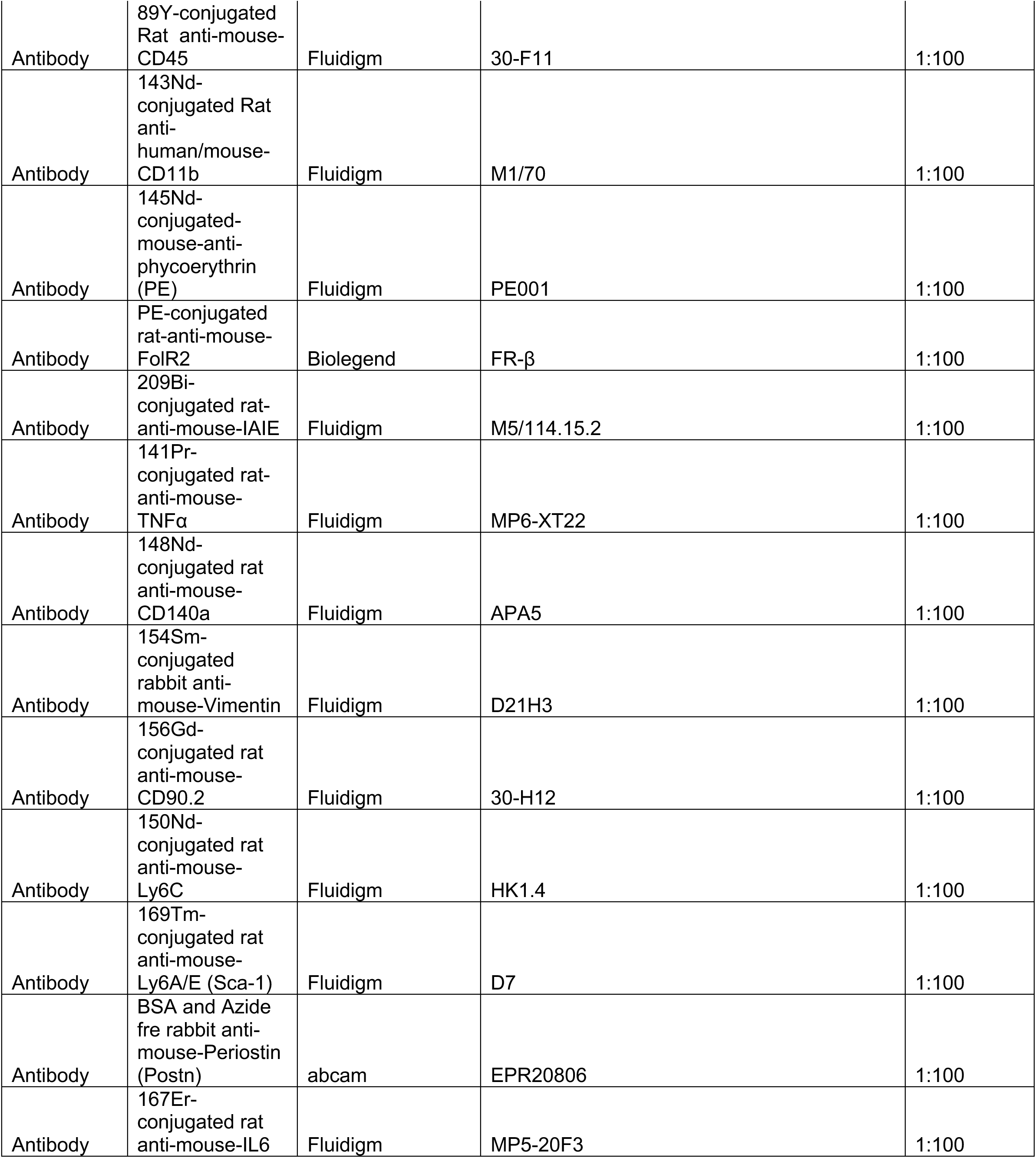
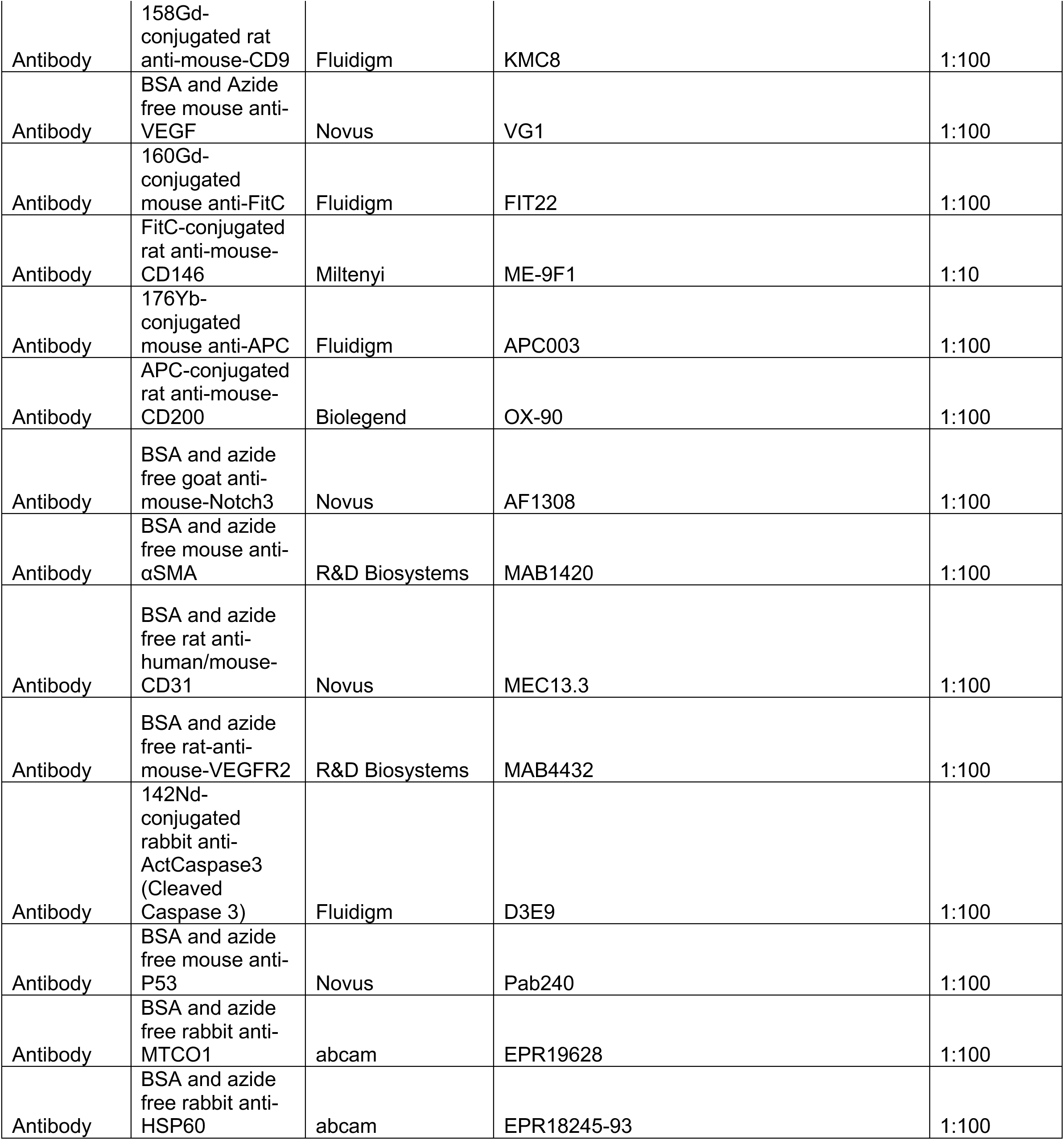
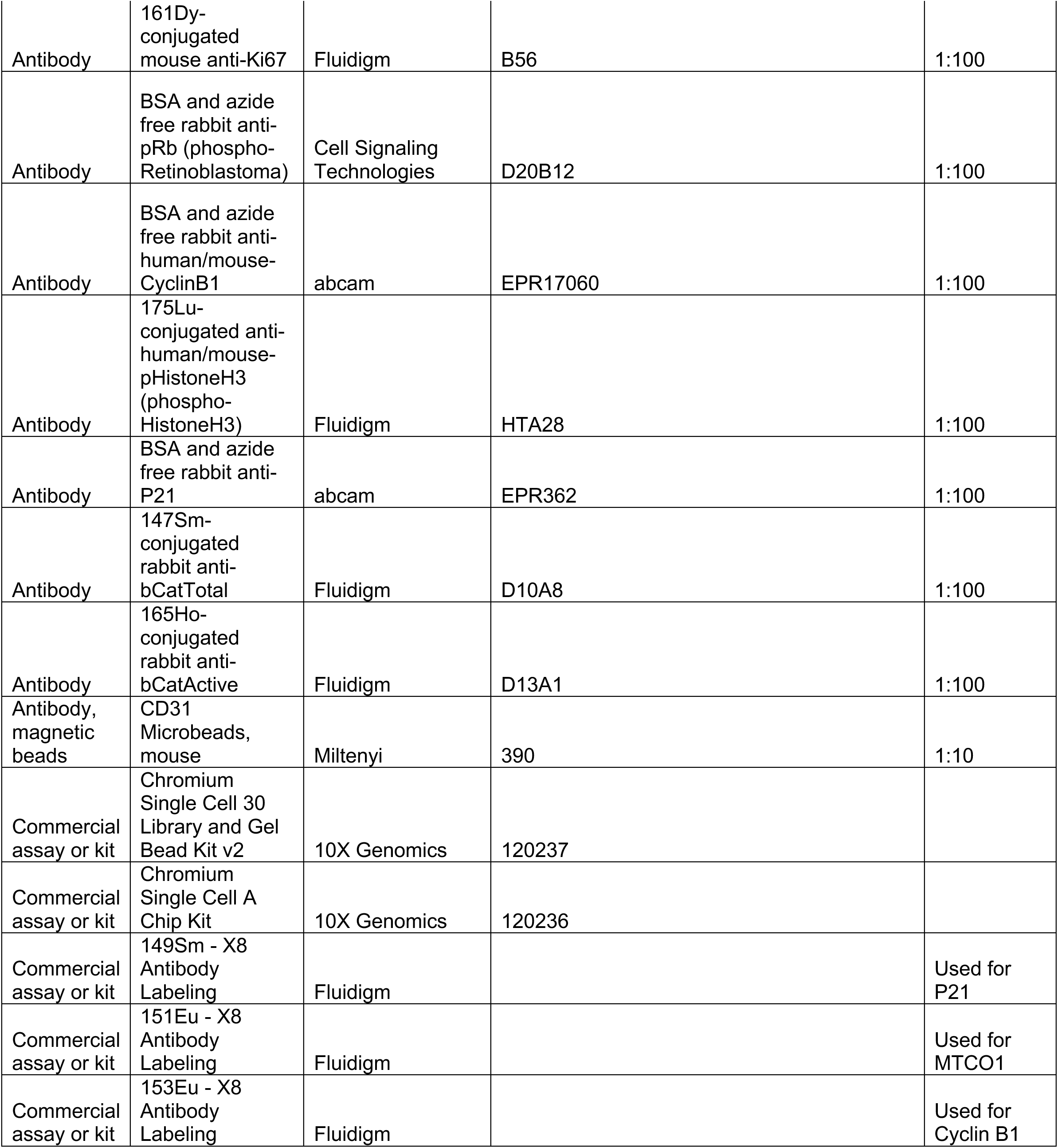
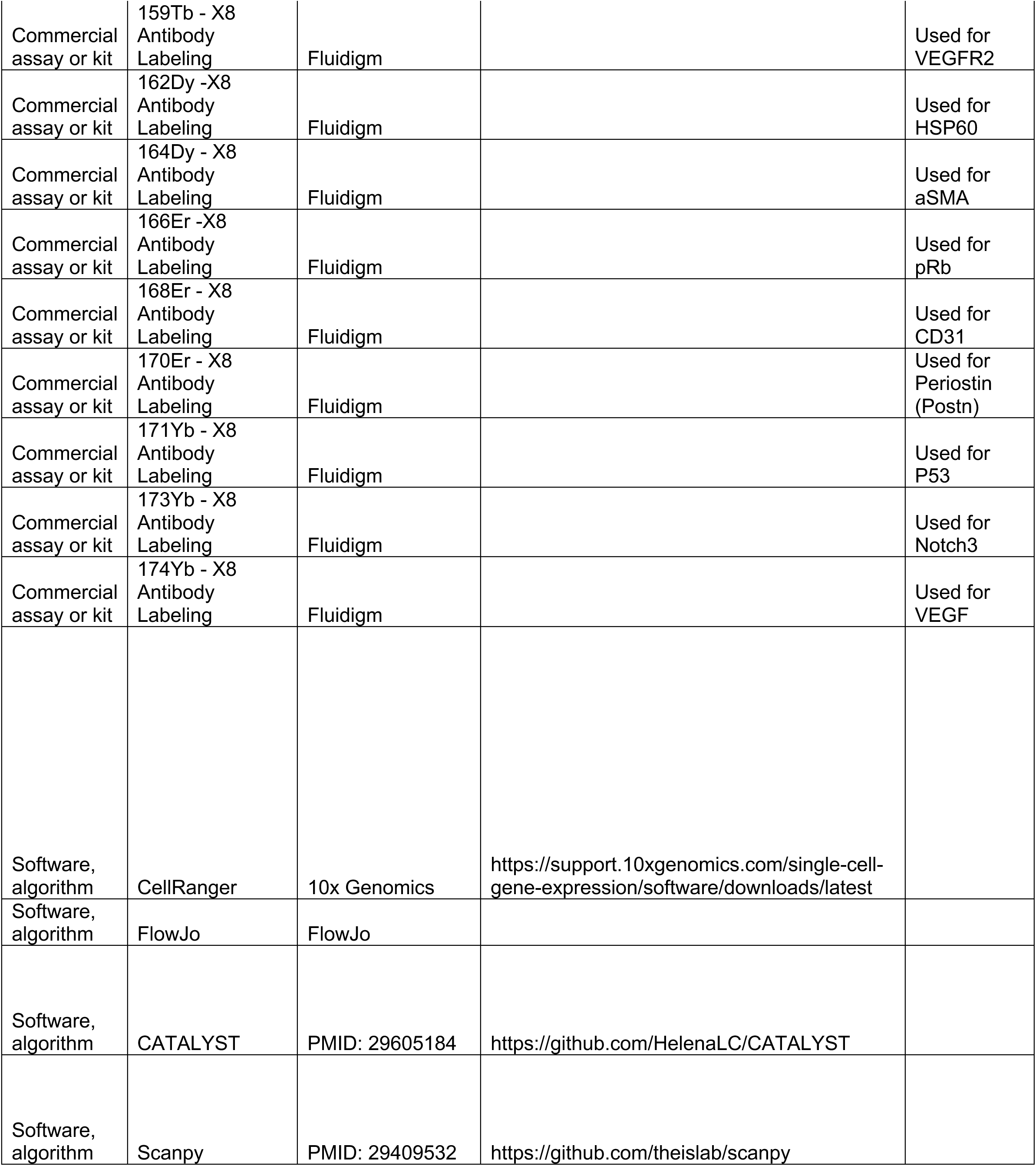
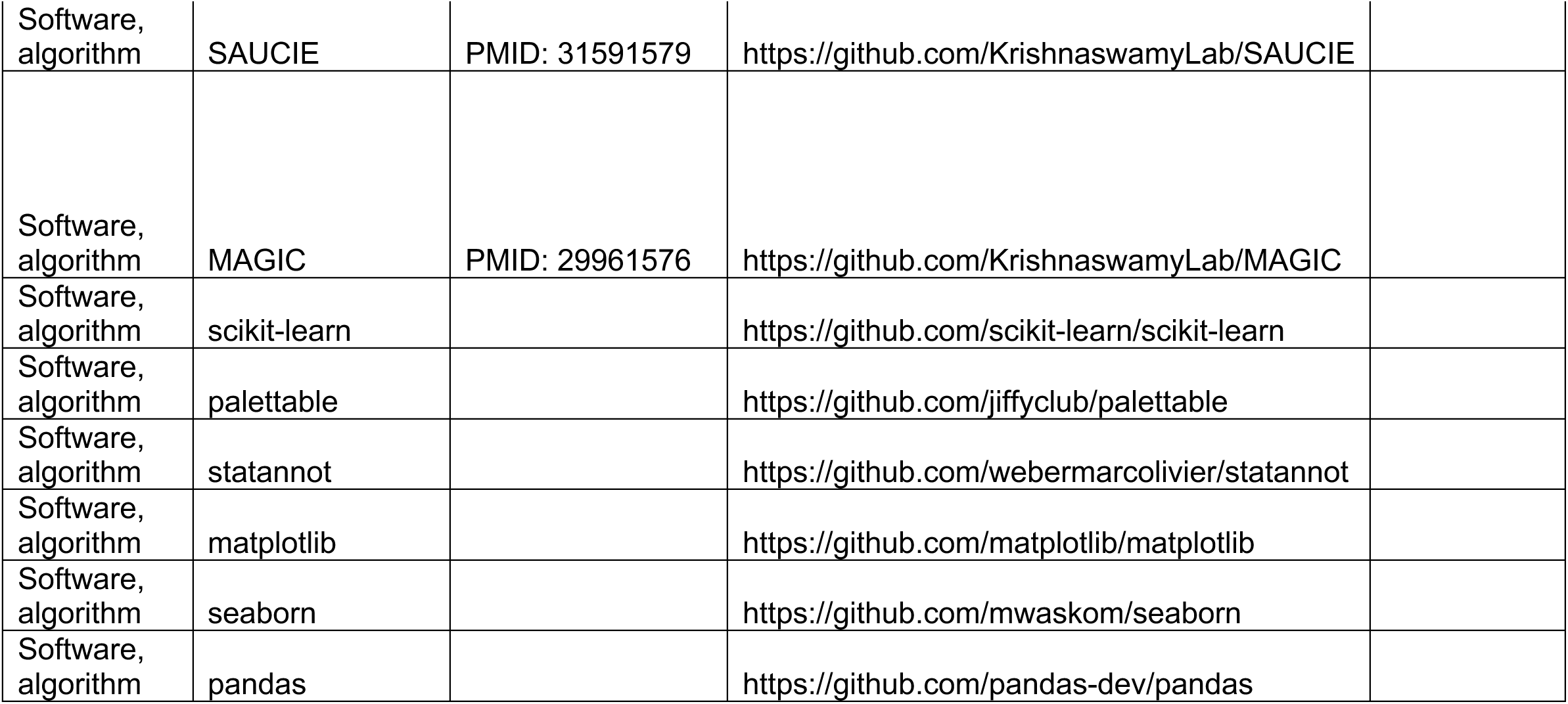

### Murine model

Mice between the ages of 8-12 weeks old were used in all experiments. Both males and females were used in the generation of our datasets. All mice were from a C57BL6/J genetic background.

### Isolation of single cell suspensions from ventricular tissue

Mice were sacrificed by CO2 inhalation followed by cervical dislocation. At necropsy, the heart was removed by cutting the heart below the atrium and residual atrial tissue was trimmed off after removal. In order to remove residual blood, ventricular tissue was washed in HBSS several times, then cut into large pieces in ice cold HBSS (without calcium or magnesium), and placed into ice cold HBSS for up to 15 minutes. Once harvest of ventricular tissue was completed, the tissue was removed from HBSS, wet with digestion buffer, and minced into fine particles with a scalpel. Similarly to Skelly, Digestion buffer consisted of collagenase IV and dispase II at a concentration of X and Y. Tissue was collected and each heart was placed into 3 mL digestion buffer, where it was then incubated in 15 mL conical tubes in a 37 degree water bath. Following 15 minutes of digestion, the digestate was triturated with a 1000 uL pipette tip, and then digested for another 15 minutes. For cardiac suspensions used in scRNAseq and CytOF, digestion was kept at 30 to 45 minutes, depending on particle size and completeness of digestion. For the pooled, overdigested reference sample, digestion was allowed to proceed for 1 hour. After digestion, cells were strained, and the strainer was washed with ice cold HBSS with 2% FBS, until a final volume of 10 mL was obtained. This suspension was washed one more time with ice cold HBSS, and then incubated in 5 mL or ACK lysis buffer for 5 minutes at room temperature to induce lysis in remaining erythrocytes. The reaction was quenched by adding an equal volumes of ice cold HBSS with 2% FBS, washed once more with HBSS with 2% FBS and then a MACS depletion or debris removal step was performed. For cell culture, the resulting suspensions were resuspended in tissue culture medium without additional processing, and supplemental growth factor or drugs were spiked in the media prior to use depending on the experiment. All centrifugation steps on cells before fixation were performed at 300 x g, unless otherwise noted; all centrifugation steps on cells after fixation were performed at 800 x g, unless otherwise noted.

### MACS depletion of CD31 positive cells

After washing cells they were incubated in anti-CD31 magnetic beads from Miltenyi according to manufacturer’s instruction. Cells were washed, then loaded onto MACS columns, and flow through was collected. The column was then discarded and the CD31 depleted flow through was then used for scRNAseq capture.

### Debris Removal and CytOF sample preparation

Cells were resuspended in 1000 µL of ice cold HBSS with 2% FBS and 300 µL of ice cold Miltenyi debris removal solution was mixed into the cell suspension. An overlay of 1000 µL was used and manufacturer’s instructions were followed for the remainder of the procedure. For the pooled reference sample, 900 µL of debris removal solution, 3100 µL of ice cold HBSS with 2% FBS was used for resuspension, and 4000 µL of ice cold HBSS with 2% FBS was used for the overlay. After overlay was added, the suspension was centrifuged at 4 degrees at 3000 x g for 10 minutes. Interphase and buffer were removed and then 2.5 mL of HBSS with 2% FBS was added to the cell suspension. The tube was gentle inverted and then centrifuged once more at 1000 x g for 10 minutes at 4 degrees, following which the supernatant was removed. Once debris was removed, cells were incubated in ice Fluidigm Cell Staining Buffer (CSB) and 0.5 µM cisplatin 198 for 15 minutes. Cells were washed once and then fixed with 1 mL 1.6% formaldehyde for 10 minutes at room temperature. After fixation, fixative was quenched by adding an equal volume of ice cold CSB and then washed once in CSB. The single cell suspensions were then resuspended in CSB with 10% DMSO and frozen in a −80 freezer until staining

### Cell culture and cell harvest

A mixture of cells from the ventricular tissue derived from 4 mice (2 male and 2 female) was seeded at a density of approximately 2 * 10^6^ cells/10 cm dish into either V media (DMEM-F12 without supplementation) or E media (DMEM-12 with 10 ng/mL hVEGFa and 10 µM Y-27632). To avoid degradation of growth factors or drugs, supplements were added prior to media use. Cells were cultured in physiological normoxia (3% O_2_) and media was changed every 3 days. On the 12^th^ day of culture, the cultures were all treated with 100 nM pharmacological grade doxorubicin dissolved in saline or vehicle. On the 14^th^ day of culture, IdU was spiked into tissue culture media to treat cells for 27 hours with a concentration of 50 µM. In order to better resolve cytokine expression, brefeldin-A (BFA) was spiked into the media 24 hours after IdU according to manufacturer’s instructions. Following 3 hours of BFA incubation, the media was removed, and cells were washed with HBSS three times prior to dissociation.

After the addition of 3 mL of TrypLE, the overconfluent cultures were dislodged from the tissue culture plastic using a cell lifter. The cell aggregates were placed into 15 mL tubes and then placed in a 37C water batch, where they were triturated every 15 minutes with a 1000 µL pipette tip. A total digestion time of 45 minutes was used to dissociate cell aggregates. After digestion, cells were strained and washed with ice cold CSB to remove digestive enzymes. The supernatant was removed following which cells were treated with 0.5 µM cisplatin 198 for 15 minutes, washed with CSB, and fixed in 1 mL 1.6% formaldehyde for 10 minutes before quenching. Cells were then washed with and frozen in CSB with 10% DMSO as previously described.

### CytOF staining procedure

On the day of staining, cells were thawed out at room temperature as well as Fluidigm 20-Plex barcodes. Cells were labeled by sample-of-origin using the barcodes according to manufacturer’s instruction. Briefly, cell suspensions were centrifuged and the supernatant freezing buffer was removed, following which samples were washed in 800 µL of 1X Barcode Perm Buffer, before being resuspended once more in 800 µL of 1X Barcode Perm Buffer. Each barcode was resuspended in 100 µL of 1X Barcode Perm Buffer and then transferred to the single cell suspensions. Samples were incubated in barcode solution for 30 minutes with gentle mixing after 15 minutes of incubation. Following centrifugation, samples were washed twice with 2 mL of CSB and resuspended in 500 µL of CSB. All the samples were then combined in a single tube for staining.

Once combined, the samples were centrifuged and the supernatant was aspirated. The cells were then resuspended in 200 µL CSB with 1:50 TruStain FcX anti-mouse CD16/32 for 10 minutes at room temperature. Following this, 200 µL of CSB with 1:50 FolR2-PE, 1:50 CD200-APC, and 1:5 CD146 -FitC were added and incubated for 30 minutes. After this incubation, cells were washed twice with 1 mL of CSB. Cells were then incubated in 350 µL of CytOF surface antibody cocktail, which included all CytOF surface markers and fluorophore secondary antibodies at a concentration of 1:100. Samples were incubated in surface staining cocktail for 30 minutes, with gentle agitation every 15 minutes. After staining, pooled samples were washed twice with CSB and then incubated with 3 mL 1X Maxpar Fix I Buffer for 10 minutes, following which they were washed twice with 6 mL of Maxpar Perm-S Buffer. The samples were then stained for 30 minutes with 350 µL of intracellular staining cocktail, which included antibodies at 1:100 concentration. Following incubation, samples were washed with 4 mL CSB twice.

Fresh fixation was performed using a fresh 1.6% formaldehyde solution from a 16% formaldehyde stock ampule. For the pooled samples, 5 mL of this solution was used to fix cells for 10 minutes at room temperature. Cells were centrifuged and the fixative was removed. Following this step, cells were incubated with Cell-ID intercalator-Ir at a concentration of 67.5 nM in 3 mL Maxpar Fix and Perm Buffer overnight. Prior to acquisition, cells were washed with CSB and then resuspended in Maxpar Cell Acquisition Solution. Maxpar Four Element Beads were put into Cell Acquisition Solution at a concentration of 10%. Cells were captured at a rate of 75-150 cells per second, to minimize doublet rate.

### Single stain bead control preparation

TotalAb compensation beads were stained in 100 µL of CSB with a concentration of 1:100 for each antibody. Beads were incubated with antibody for 1 hour at room temperature and then washed twice with CSB. The beads were then pooled and fixed with 1.6% formaldehyde for 1 hour at room temperature. They were then washed twice with CSB and kept in the fridge overnight before the CytOF run.

### Capture and sequencing with 10x Genomics

10x Genomics is a commercially available UMI and droplet-based single cell sequencing capture method. Manufacturer’s instructions were followed for this experiment. Libraries were generated following capture and were sequenced on an Illumina NextSeq 500.

### Processing of 10x Genomics scRNAseq data

Using 10x Genomics CellRanger 3.0.2, BCL files were processed to fastqs using CellRanger mkfastq. Fastq files were aligned to the mm10 2.1.0 reference genome. Following this, a count matrix was generated using CellRanger count.

### Reprocessing of previously reported 10x Genomics scRNAseq data

Fastqs from Skelly and Farbehi were downloaded and recounted using 10x Genomics CellRanger 3.0.2 count. Both of the Skelly datasets were used in the study. From Farbehi, only ‘Sham TIP Day 7’, ‘Sham GFP Day 7’, ‘MI TIP Day 3’, ‘MI TIP Day 7’, and ‘MI GFP Day 7’ (MI – myocardial infarction) were able to be recounted with Cellranger 3.0.2. Fastqs that produced errors when recounted which terminated the count program were excluded from the study.

### Processing of CytOF data

Raw FCS files were retrieved from the computer interfaced with the Helios CytOF. Using CATALYST, compensation bead and barcoded experiment data were normalized. Compensation beads were used to define a spillover matrix, which was then applied to the barcoded experiment. The barcoded experiment was then debarcoded, yielding 11 separate FCS files. The resulting FCS files were then taken into FlowJo and Fluidigm Guassian channels were used to further filter out doublets. Following this, the FCS files were opened with flowCore and unrandomized (using a ceiling function) hyperbolic arcsin (with a factor of 5) transformed protein expression matrices were extracted from each FCS for secondary analysis.

### Quality control of CytOF data

CytOF data was imported into Scanpy (1.4.5) where quality control cut-offs for *in vivo* CytOF were set a minimum of 30 transformed counts, a maximum of 60 transformed counts, a minimum of 15 detected proteins, and a maximum of 25 detected proteins. Cut-offs for *in vitro* CytOF were set at a minimum of 50 transformed counts, a maximum of 85 transformed counts, a minimum of 20 detected proteins, and a maximum of 32 detected proteins. For *in vivo* data, from a total of 436,247 cells, 343,245 cells passed QC. For *in vitro* data, from a total of 168,459 cells, 125,109 cells passed QC. Cut-offs were determined by inspecting count as well as protein distributions using violin plots and intended to trim outliers, in order to prevent SAUCIE from overfitting on low quality cells.

### Quality control of scRNAseq data

For the initial analysis of the CD31 depleted (Iskra) dataset to identify robust CytOF markers, we set a cut-off for cells with at least 200 genes detected, at most 4000 genes, genes detected in at least 3 cells, and a percent mitochondrial content of less than 17.5% genes detected. Top 2500 highly variable genes were selected to generate 30 principal components, which were then used to build the nearest neighbor graph with a k of 30. Leiden community detection was used and a UMAP embedding was generated with PAGA to initialize positions.

scRNAseq datasets were imported into Scanpy where quality control cut-offs were set a minimum of 750 genes, a maximum of 3750 genes, a minimum of 1250 counts, a maximum of 12500 counts, and a mitochondrial content of less than 17.5% of total genes detected. Genes expressed in less than 250 cells were excluded from the analysis. Data was log plus one transformed and then processed with MAGIC using 30 principal components (otherwise, default scanpy settings were used) prior to integration in SAUCIE.

### Data integration with SAUCIE

Post-quality control data was put into a data matrix and batch labels were set according to data modality. To ‘convert’ CytOF data to scRNAseq data, Ly6AE/Sca-1 expression was set to the sum of Ly6a and Ly6e expression, IdU was set to Mki67 expression, b-Catenin-Active expression was set to Dkk3 expression, pHistoneH3 expression was set to Birc5 expression, pRb expression was set to Rb1 expression, Active Caspase3 expression was set to Casp3 expression. All scRNAseq data was labeled as batch 0, whereas *in vivo* CytOF was set as batch 1, and *in vitro* CytOF was set as batch 2. With these settings, all CytOF protein data are transformed to scRNAseq gene expression. Data from scRNAseq and *in vitro* CytOF were sampled with replacement to ensure equivalent representation in the training dataset with respect to *in vivo* CytOF data. SAUCIE was run deterministically on a CPU with a random seed of 42 for numpy, random, and tensorflow.

A test-train split of 20% and 80%, respectively, was used to ensure that SAUCIE was not overfitting on the training data. For data integration, SAUCIE was run with layers set to [1024, 512, 256, 2], a lambda b equal to 0.00625, and trained for 2,000 steps. These settings were determined empirically and chosen to allow for the longest training session in terms of steps before overfitting, as determined by inspected the information bottleneck layer of SAUCIE to determine a meaningful distribution of genes where key cell identities (endothelium, fibroblasts, leukocytes, and pericytes) were able to be resolved. For clustering the integrated data, SAUCIE was run with layers set to [512, 256, 128, 2] with a lambda c equal to 0.05 and a lambda d equal to 0.10. Training was performed for 8,000 steps and 9 clusters were detected, which were then merged according to gene expression and localization on the shared latent space representation. SAUCIE-determined labels were then transferred to individual CytOF or scRNAseq datasets during secondary analysis.

### Secondary analysis of CytOF data

Using Scanpy, post quality control cells were labeled with SAUCIE labels and counts were normalized to 50 hyperbolic arcsin transformed counts per cell, for both *in vivo* and *in vitro* CytOF. Each of the 6 non-reference *in vivo* samples were subsampled randomly such that 7,500 cells from each were represented in the analysis. For *in vitro* samples, 11,250 cells were randomly sampled from each condition. On these subsets of cells, all markers were used for constructing the nearest neighbor graph, where 30 nearest neighbors were selected for the graph. Louvain clustering was then performed with a resolution of 1. PAGA and UMAP were computed with default scanpy settings and PAGA was used to initialize the UMAP embedding.

## Acknowledgements

Special thanks to the UTSA Genomics core for their assistance with the 10X Genomics platform as well as the Greehey Children’s Cancer Research Institute Genome Sequencing Facility.

## Figures and Tables

**Figure 2 Supplement 1.**
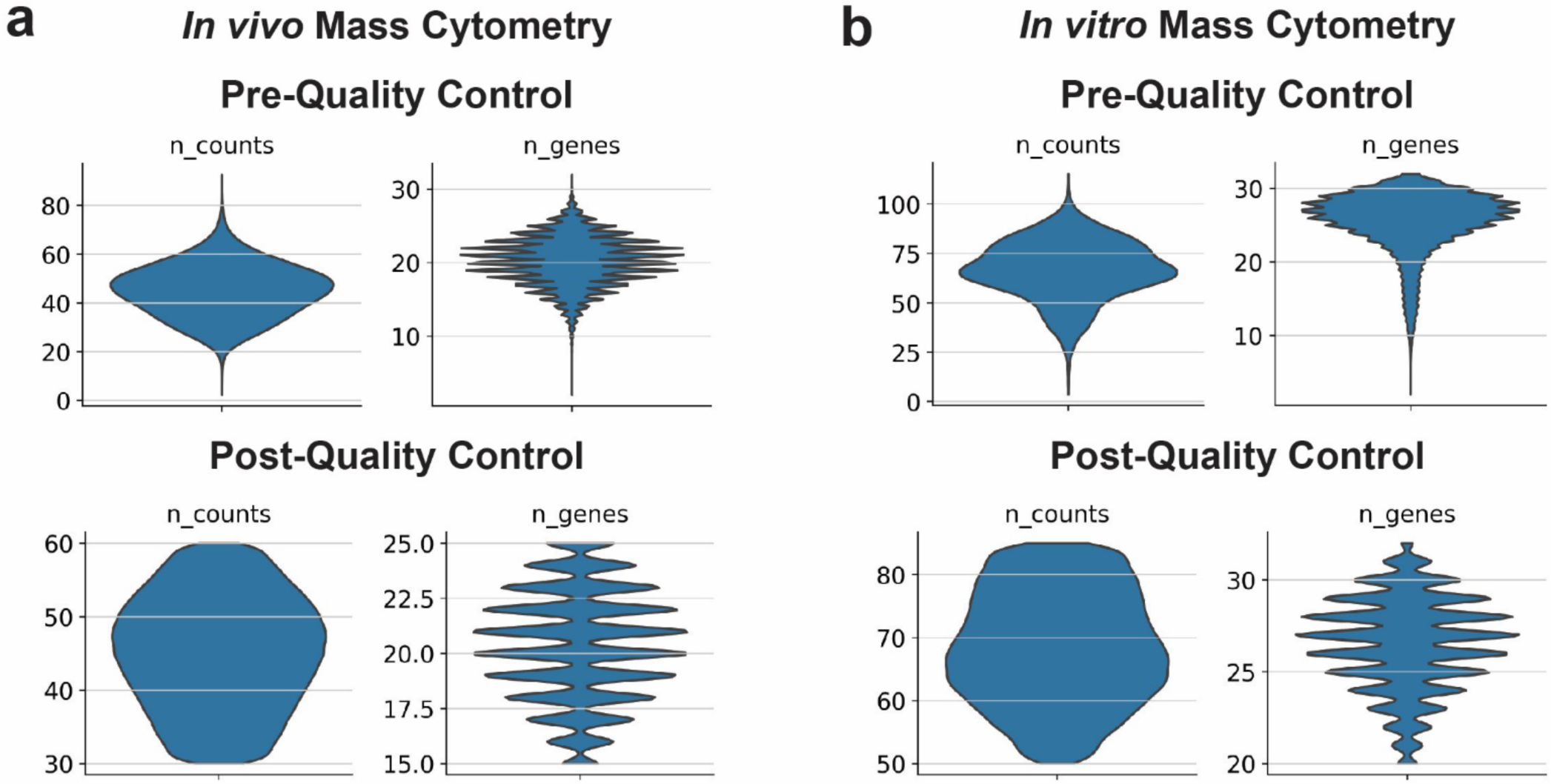
Quality control information and cut-offs for CytOF data. **a)** Per cell distribution of hyperbolic arcsin factor 5 transformed counts and number of genes expressed in each cell as found i*n vivo* CytOF data.

**Figure 6 Supplement 1.**
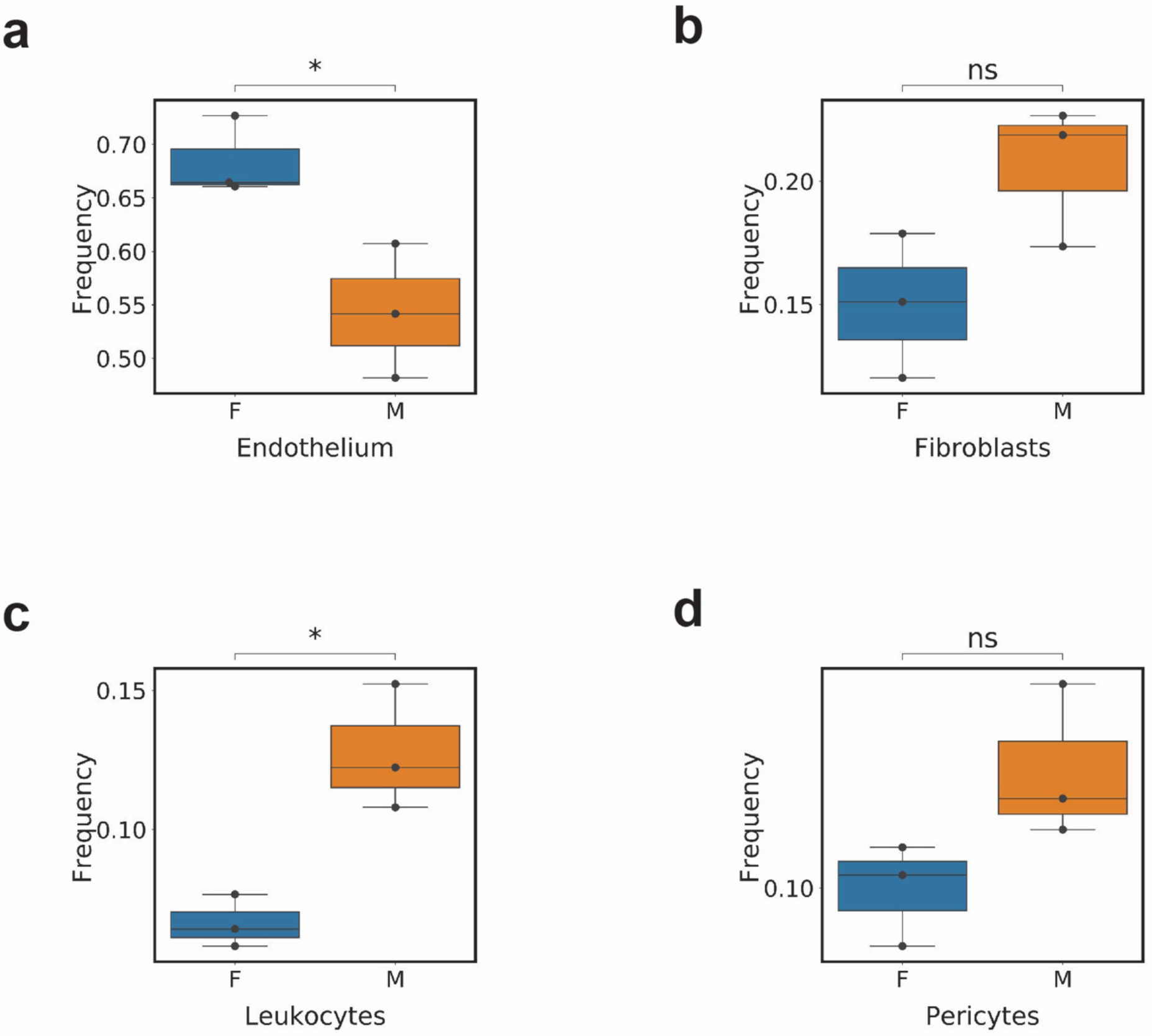
Comparative frequency of major populations by mouse sex detected in the CytOF dataset. **a)** Boxplot of endothelium frequency in female mice versus male mice. **b)** Boxplot of fibroblasts frequency in female mice versus male mice. **c)** Boxplot of leukocytes frequency in female mice versus male mice. **d)** Boxplot of pericytes frequency in female mice versus male mice. * signifies a p-value of less than 0.05.

## Bibliography

Ackers-Johnson Matthew, Li Peter Yiqing, Holmes Andrew P., O’Brien Sian-Marie, Pavlovic Davor, & Foo Roger S. (2016). A Simplified, Langendorff-Free Method for Concomitant Isolation of Viable Cardiac Myocytes and Nonmyocytes From the Adult Mouse Heart. Circulation Research, 119(8), 909–920. https://doi.org/10.1161/CIRCRESAHA.116.309202

Amodio, M., van Dijk, D., Srinivasan, K., Chen, W. S., Mohsen, H., Moon, K. R., Campbell, A., Zhao, Y., Wang, X., Venkataswamy, M., Desai, A., Ravi, V., Kumar, P., Montgomery, R., Wolf, G., & Krishnaswamy, S. (2019). Exploring single-cell data with deep multitasking neural networks. Nature Methods, 16(11), 1139–1145. https://doi.org/10.1038/s41592-019-0576-7

Chevrier, S., Crowell, H. L., Zanotelli, V. R. T., Engler, S., Robinson, M. D., & Bodenmiller, B. (2018). Compensation of Signal Spillover in Suspension and Imaging Mass Cytometry. Cell Systems, 6(5), 612–620.e5. https://doi.org/10.1016/j.cels.2018.02.010

Cieslik, K. A., Trial, J., & Entman, M. L. (2015). Mesenchymal stem cell-derived inflammatory fibroblasts promote monocyte transition into myeloid fibroblasts via an IL-6-dependent mechanism in the aging mouse heart. The FASEB Journal, 29(8), 3160–3170. https://doi.org/10.1096/fj.14-268136

Dijk, D. van, Sharma, R., Nainys, J., Yim, K., Kathail, P., Carr, A. J., Burdziak, C., Moon, K. R., Chaffer, C. L., Pattabiraman, D., Bierie, B., Mazutis, L., Wolf, G., Krishnaswamy, S., & Pe’er, D. (2018). Recovering Gene Interactions from Single-Cell Data Using Data Diffusion. Cell, 174(3), 716–729.e27. https://doi.org/10.1016/j.cell.2018.05.061

Farbehi, N., Patrick, R., Dorison, A., Xaymardan, M., Janbandhu, V., Wystub-Lis, K., Ho, J. W., Nordon, R. E., & Harvey, R. P. (2019). Single-cell expression profiling reveals dynamic flux of cardiac stromal, vascular and immune cells in health and injury. ELife, 8, e43882. https://doi.org/10.7554/eLife.43882

He, L., Huang, X., Kanisicak, O., Li, Y., Wang, Y., Li, Y., Pu, W., Liu, Q., Zhang, H., Tian, X., Zhao, H., Liu, X., Zhang, S., Nie, Y., Hu, S., Miao, X., Wang, Q.-D., Wang, F., Chen, T., … Zhou, B. (2017). Preexisting endothelial cells mediate cardiac neovascularization after injury. The Journal of Clinical Investigation, 127(8), 2968–2981. https://doi.org/10.1172/JCI93868

Hulsmans, M., Clauss, S., Xiao, L., Aguirre, A. D., King, K. R., Hanley, A., Hucker, W. J., Wülfers, E. M., Seemann, G., Courties, G., Iwamoto, Y., Sun, Y., Savol, A. J., Sager, H. B., Lavine, K. J., Fishbein, G. A., Capen, D. E., Da Silva, N., Miquerol, L., … Nahrendorf, M. (2017). Macrophages Facilitate Electrical Conduction in the Heart. Cell, 169(3), 510–522.e20. https://doi.org/10.1016/j.cell.2017.03.050

Janssen, W. J., Bratton, D. L., Jakubzick, C. V., & Henson, P. M. (2016). Myeloid cell turnover and clearance. Microbiology Spectrum, 4(6). https://doi.org/10.1128/microbiolspec.MCHD-0005-2015

Joo, H. J., Choi, D.-K., Lim, J. S., Park, J.-S., Lee, S.-H., Song, S., Shin, J. H., Lim, D.-S., Kim, I., Hwang, K.-C., & Koh, G. Y. (2012). ROCK suppression promotes differentiation and expansion of endothelial cells from embryonic stem cell–derived Flk1+ mesodermal precursor cells. Blood, 120(13), 2733–2744. https://doi.org/10.1182/blood-2012-04-421610

Kluge, M. A., Fetterman, J. L., & Vita, J. A. (2013). Mitochondria and Endothelial Function. Circulation Research, 112(8), 1171–1188. https://doi.org/10.1161/CIRCRESAHA.111.300233

Kretzschmar, K., Post, Y., Bannier-Hélaouët, M., Mattiotti, A., Drost, J., Basak, O., Li, V. S. W., Born, M. van den, Gunst, Q. D., Versteeg, D., Kooijman, L., Elst, S. van der, Es, J. H. van, Rooij, E. van, Hoff, M. J. B. van den, & Clevers, H. (2018). Profiling proliferative cells and their progeny in damaged murine hearts. Proceedings of the National Academy of Sciences, 115(52), E12245–E12254. https://doi.org/10.1073/pnas.1805829115

Maliken Bryan D., Kanisicak Onur, Karch Jason, Khalil Hadi, Fu Xing, Boyer Justin G., Prasad Vikram, Zheng Yi, & Molkentin Jeffery D. (2018). Gata4-Dependent Differentiation of c-Kit+–Derived Endothelial Cells Underlies Artefactual Cardiomyocyte Regeneration in the Heart. Circulation, 138(10), 1012–1024. https://doi.org/10.1161/CIRCULATIONAHA.118.033703

Omelchenko, T., Vasiliev, J. M., Gelfand, I. M., Feder, H. H., & Bonder, E. M. (2002). Mechanisms of polarization of the shape of fibroblasts and epitheliocytes: Separation of the roles of microtubules and Rho-dependent actin-myosin contractility. Proceedings of the National Academy of Sciences, 99(16), 10452–10457. https://doi.org/10.1073/pnas.152339899

Pinto, A. R., Ilinykh, A., Ivey, M. J., Kuwabara, J. T., D’Antoni, M. L., Debuque, R., Chandran, A., Wang, L., Arora, K., Rosenthal, N. A., & Tallquist, M. D. (2016). Revisiting Cardiac Cellular Composition. Circulation Research, 118(3), 400–409. https://doi.org/10.1161/CIRCRESAHA.115.307778

Skelly, D. A., Squiers, G. T., McLellan, M. A., Bolisetty, M. T., Robson, P., Rosenthal, N. A., & Pinto, A. R. (2018). Single-Cell Transcriptional Profiling Reveals Cellular Diversity and Intercommunication in the Mouse Heart. Cell Reports, 22(3), 600–610. https://doi.org/10.1016/j.celrep.2017.12.072

Trial, J., Heredia, C. P., Taffet, G. E., Entman, M. L., & Cieslik, K. A. (2017). Dissecting the role of myeloid and mesenchymal fibroblasts in age-dependent cardiac fibrosis. Basic Research in Cardiology, 112(4), 34. https://doi.org/10.1007/s00395-017-0623-4

Zunder, E. R., Finck, R., Behbehani, G. K., Amir, E. D., Krishnaswamy, S., Gonzalez, V. D., Lorang, C. G., Bjornson, Z., Spitzer, M. H., Bodenmiller, B., Fantl, W. J., Pe’er, D., & Nolan, G. P. (2015). Palladium-based Mass-Tag Cell Barcoding with a Doublet-Filtering Scheme and Single Cell Deconvolution Algorithm. Nature Protocols, 10(2), 316–333. https://doi.org/10.1038/nprot.2015.020

